# A genome-wide-level insight into the HSF gene family of *Rhodomyrtus tomentosa* and the functional divergence of *RtHSFA2a* and *RtHSFA2b* in thermal adaption

**DOI:** 10.1101/2024.10.14.618362

**Authors:** Hui-Guang Li, Ling Yang, Yujie Fang, Gui Wang, Shanwu Lyu, Shulin Deng

**Affiliations:** Key Laboratory of National Forestry and Grassland Administration on Plant Conservation and Utilization in Southern China, Guangdong Provincial Key Laboratory of Applied Botany, and Xiaoliang Research Station for Tropical Coastal Ecosystems, South China Botanical Garden, Chinese Academy of Sciences, Guangzhou 510650, China; College of Life Sciences, Gannan Normal University, Ganzhou 341000, China; University of Chinese Academy of Sciences, Beijing 100049, China

**Author notes:** Author for correspondence: Shulin Deng.

**Keywords:** *Rhodomyrtus tomentosa*, heat shock transcription factor, HSFA2, thermal adaption, functional divergence

## Abstract

Heat shock transcription factor (HSF) is one of the most important regulatory elements in plant development and stress response. *Rhohomyrtus tomentosa* has many advantages in adapting to high temperature and humidity climates, whereas the inherence has barely been elucidated. In this study, we aimed to characterize the HSF family and investigate the thermal adaption mechanisms of *R. tomentosa*. We identified 25 HSF genes in the *R. tomentosa* genome. They could be classified into three classes: HSFA, HSFB, and HSFC. Gene duplication event is a major motivation for the expansion of the *RtHSF* gene family. Most of the genes in the same subclass share similar conserved motifs and gene structures. The *cis-acting* elements of the promoter regions of *RtHSF* genes are related to development, phytohormone signaling, and stress responses, and they vary among the genes even in the same subclass, resulting in different expression patterns. Especially, there exists subfunctionalization in the *RtHSFA2* subfamily in responding to various abiotic stresses, *viz. RtHSFA2a* is sensitive to drought, salt, and cold stresses, whilst *RtHSFA2b* is mainly induced by heat stress. We further prove that *RtHSFA2b* might be of more importance in *R. tomentosa* thermotolerance, for *Arabidopsis* with overexpressed *RtHSFA2b* outperformed those with *RtHSFA2a* under heat stress, and *RtHSFA2b* has much higher transcription activity than *RtHSFA2a* in regulating certain heat shock response (HSR) genes. *RtHSFA2a* plays a role in transactivating *RtHSFA2b*. All these results provide a general prospect of the *RtHSF* gene family and enclose a basal thermal adaption mechanism of *R. tomentosa*.

## 1. Introduction

Plants are the most important organisms in softening and thriving the primitive Earth by harvesting solar energy, consuming atmospheric CO_2_, releasing O_2,_ and providing organic matter therefore sustaining the global ecosystem. When migrating from the sea to the land, they require various morphology and metabolism alterations to conquer the Pangaea (Kenrick and Crane, 1997; Schreiber et al., 2022). One of the challenges plants have to subdue is the fickle temperature due to the unstable crustal movement and thin ozone (Biggin et al., 2015; Doglioni et al., 2016; Schreiber et al., 2022), which poses great threats to the survival of terrestrial biota. Even in this day, ground surface temperature is still one of the factors limiting worldwide arable land utilization and restricting crop productivity (Lobell et al., 2011; Zhang et al., 2022a), and the acceleration of global warming driven by anthropogenic carbon emission from fossil fuels impacts agri-forestry yield and biodiversity on a larger scale (Pecl et al., 2017; Zandalinas et al., 2021; Zhang et al., 2019). However, the unprecedented increases in human population growth and demands are exacerbating the pace of climate change, usually beyond our expectations (Flores et al., 2024; Mo et al., 2023).

Extreme high temperature brings deleterious influences to plants, including accumulation of disrupted protein and burst of reactive oxygen species (ROS), leading to growth retardation or even mortality (Kan et al., 2021, 2023). Accordingly, plants employ various regulatory mechanisms to offset the detrimental effects of temperature fluctuations, such as ROS scavenging, morphology alteration, and renaturation or elimination of denatured proteins (Qin et al., 2022; Zhang et al., 2019). Among these transcriptional regulations or metabolism modifications, heat shock transcription factor (HSF) is one of the most crucial gene families that functions not only in stress response but also in plant morphogenesis and propagation. The structure of the HSF family is highly conserved among plants: a highly helix-turn-helix (HTH) DNA-binding domain (DBD) at the N-terminus, followed by a bipartite heptad repeat pattern (HR-A/B region) oligomerization domain (OD), and nuclear localization signaling domain (NLS) (Harrison et al., 1994; Scharf et al., 2012; Xie et al., 2023). According to the length of the flexible linker region between DBD and HR-A/B regions and the number of amino acid residues inserted into the HR-A/B regions, HSFs are classified into three classes, *viz.* HSFA, B, and C (Kotak et al., 2004; Nover et al., 2001). Class A HSFs possess activator motifs or aromatic and hydrophobic amino acid residues (AHAs) close to the NES at the C-terminal region, while class B HSFs (except HSFB5) contain a characteristic tetrapeptide (-LFGV-) repressor domain (RD) instead for transcription suppression (Andrási et al., 2021; Guo et al., 2016; Nover et al., 2001).

HSF family modulates plant development and stress response by regulating a set of genes. According to current reports, HSFA1 acts as a master regulator of plant heat stress response (HSR) (Liu et al., 2011; Wang et al., 2023). HSFA2 and HSFA3 are induced by heat stress under the persuasion of HSFA1 and function vitally by regulating heat stress response genes like heat shock proteins (HSP) or by modifying growth and metabolism (Andrási et al., 2021; Friedrich et al., 2021; Liu et al., 2023; Wang et al., 2023). HSFA4 genes act as activators of heat shock and desiccation response genes, while HSFA5 functions as a repressor of HSFA4 activity (Baniwal et al., 2007; Lang et al., 2017). In apple, MdHSFA8a is involved in flavonoid synthesis and associated with abscisic acid-induced stomatal closure (Wang et al., 2020). HSFA9 takes part in late embryogenesis and early photomorphogenesis (Prieto-Dapena et al., 2017), UV-B light responses (Carranco et al., 2022), and severe dehydration response (Prieto-Dapena et al., 2008). The most common cognition about the B class of HSFs is acting as respressors (Ohama et al., 2017). Recent research in maize demonstrates that maize HSFB2a negatively modulates heat stress tolerance whereby transcriptionally repressing *ZmHSF4* and *ZmCesA2* (Li et al., 2024). Other studies reveal that HSFB1 and HSFB2b suppress heat shock response genes under non-heat-stress conditions while are necessary for acquired thermotolerance (Ikeda et al., 2011), and another study proves that HSFB1 is induced by heat and confers thermotolerance to grapevine (Chen et al., 2023b). Class C HSFs are still underexamined. A recent study revealed that lily HSFC2 can interact with other HSFs and play an active role in the general balance and maintenance of heat stress response (Wu et al., 2024).

*Rhodomyrtus tomentosa* (Ait.) Hassk, a shrub of the Mytaceae family, thrives in the hilly area of Southern China, Malaysia, Indonesia, Philippines, and India (Wang et al., 2022b). *R. tomentosa* is commonly known as a wild healthy fruit that could be eaten fresh or made into juice, jam, and beverages (Wang et al., 2022a; Yang et al., 2023). In China, Vietnam, and Malaysia, the whole plant of *R. tomentosa* is used as a traditional herbal medicine to treat diarrhea, chronic dysentery, and stomachaches (Zhao et al., 2020; Zhuang et al., 2017). *R. tomentosa* is rich in phenolic compounds and possesses multiple pharmacological effects in antibacterial activity (Srisuwan et al., 2018) or in preventing chronic diseases like hypercholesterolemia and atherosclerosis (Sinaga et al., 2021). Profit from the large inflorescence and long flowering period, *R. tomentosa* nowadays is favored in gardening. Additionally, *R. tomentosa* favors sunshine and is underdemanding about soil conditions (Lai et al., 2015), making it a superb pioneer tree species for afforestation or rehabilitation in the waste mountain. However, despite the immense potential *R. tomentosa* possesses in food production, pharmaceuticals industry, horticulture, and ecological remediation, it is still lack of exploitation and utilization. Considering the habitat of *R. tomentosa*, which is always combined with acid soil, barren, high radiation, and ambient temperature (Bhat et al., 2010; Kochian et al., 2015), investigating the hereditary basis of *R. tomentosa* is of vast value in demonstrating the mechanisms of *R. tomentosa* in adapting to its hostile environment, as well as providing gene resource and strategy for tree breeding.

Recently, we assembled a high-quality whole genome of *R. tomentosa* with annotated chromosomal level details (Yang et al., 2024), opening up a wealth of doors to investigate the mechanisms including secondary metabolites synthesis or stress response. In this study, we aim to identify the HSF family of *R. tomentosa,* investigate their structural features and evolutionary relationships, and reveal their hidden roles in heat stress response. Our results showed that the *R. tomentosa* genome encodes 25 HSF members with a particular propensity of functional differentiations in adapting to various stresses. This study provided a comprehensive perception of the *RtHSF* family which is not only valuable in illustrating the mechanism underlying plant adapting to tropic and subtropic climates but also in gene modification for stress-tolerant plants.

## 2. Materials and methods

### 2.1 Data accession and genome-wide identification of *Rt*HSF genes

The genome data of *Arabidopsis thaliana*, *Eucalyptus grandis*, *Oryza sativa*, and *Populus trichocarpa* were derived from Phytozome (Goodstein et al., 2012). The gene ids of *AtHSF*, *EgHSF*, *OsHSF*, and *PtHSF* gene families were obtained from previous publications (Mittal et al., 2009; Yuan et al., 2022; Zhang et al., 2015) and PlantTFDB (https://planttfdb.gao-lab.org/, last accessed on 2 March 2024).

The genome data of *R. tomentosa* we published was used to identify the HSF family using the hidden Markov model (PF00447), obtained from InterPro (https://www.ebi.ac.uk/interpro/ac.uk/, last accessed on 1 March 2024). The function “hmmsearch” of the HMMER software (version 3.4, http://hmmer.org/, last accessed on 2 March 2024) was executed to screen the total protein sequences of *R. tomentosa* and the identified genes with e-values less than or equal to 1E-5 were considered putative candidates. The HSF classic motifs of all these putative sequences were then verified by the SMART program (http://smart.embl-heidelberg.de/, last accessed on 1 March 2024) and HEATSTER platform (https://applbio.biologie.uni-frankfurt.de/hsf/heatster/, last accessed on 1 March 2024).

2.2 Phylogenetic analysis and classification of *R. tomentosa*

The protein sequences of HSF genes of *R. tomentosa*, *A. thaliana*, *and E. grandis* were subjected to multiple sequence alignment using the Muscle alignment tools (version 11.0.13, https://www.drive5.com/muscle/, last accessed on 2 March 2024). A phylogenetic tree was then constructed with the maximum likelihood (ML) method using RAxML-NG software (Kozlov et al., 2019) (version 1.2.1, https://github.com/amkozlov/raxml-ng, last accessed on 2 March 2024) with a bootstrap value at 1000. The output of RAxML-NG was visualized by the ITOL online server (https://itol.embl.de/, last accessed on 2 March 2024).

### 2.3 Primary sequences analysis of *RtHSF* members

We used TBtools version 2.0.30 (Chen et al., 2023a) (https://github.com/CJ-Chen/TBtools-II/, last accessed on 4 March 2024) to plot the chromosomal positions of identified *RtHSF* genes. The Gene Structure Display Server (GSDS, http://gsds.gao-lab.org/, last accessed on 3 March 2024) was used to illustrate the gene structures. The tool ProtParam of Expasy (https://web.expasy.org/protparam/, last accessed on 3 March 2024) was used to predict the molecular weight (MW) and isoelectric point. The subcellular localization of *RtHSF* proteins was predicted by the Plant-mPLoc (http://www.csbio.sjtu.edu.cn/bioinf/plant-multi/, last accessed on 3 March 2024). To detect the conserved motifs of *RtHSF* proteins, we applied the MEME suite (https://meme-suite.org/meme/tools/meme, last accessed on 2 March 2024) to analyze the protein sequences with the parameters configured to “motif width = 6 to 200” and “Number of motifs = 10”. The promoter sequences of *RtHSF* genes (2 kb upstream of the open reading frame) were extracted from the *R. tomentosa* genome and the *cis-acting* elements were predicted by the PlantCARE database (https://bioinformatics.psb.ugent.be/webtools/plantcare/html/, last accessed on 2 March 2024) and mapped by TBtools.

### 2.4 Chromosomal distribution, synteny, and gene duplication analyses

The localization of *RtHSF* genes on *R. tomentosa* chromosomes was mapped using TBtools. Synteny analysis among *RtHSF* genes was also executed by TBtools. The Python version of MCScanX (JCVI) (Wang et al., 2012) was used to identify the synteny blocks among *R. tomentosa*, *A. thaliana*, *E. grandis*, and *P.trichocarpa* with default parameters. Gene duplication analysis was performed using DupGen_finder (Qiao et al., 2019; Yang et al., 2020) with default parameters. To assess the selection pressure, we estimated the synonymous substitution (Ks) and nonsynonymous substitution (Ka) values as well as the Ka/Ks rate of the duplicated gene pairs by TBtools (Wang et al., 2017b).

### 2.5 Plant growth conditions and stresses treatment

*A. thaliana* Columbia-0 (Col-0) ecotype was used in this research. Seeds were surface sterilized with 75% ethanol for 15 min followed by rinsing with 100% ethanol and then paved on 1/2 MS medium containing 0.54% agar and 1% sucrose and vernalized at 4 °C in darkness for 2 days. After that, they were transferred to a phytotron with a 16-h-light / 8-h-dark regime at 22 °C. Six-day-old seedlings were planted in soil and grew in the same conditions.

*R. tomentosa* seeds were collected from the South China National Botanical Garden (Guangzhou, Guangdong Province, China) and germinated in a phytotron in soil supplied with a 16/8 h (light/dark) photoperiod at 25 °C. Sprouted seedlings were then transplanted into pots (10 cm × 10 cm, nine plantlets in each pot) and maintained under the same conditions.

Comparable seedlings of *R. tomentosa* grew in soil for three months were selected for stress treatments. For heat stress, plantlets were exposed to 45 °C without previous accumulation for 0 h, 1 h, and 6 h. Leaves were collected after that. For cold treatment, seedlings were transferred to 4 °C, and leaves were collected at 0 h, 3 h, 6 h, 12 h, and 24 h. For drought treatment, twelve comparable pots of *R. tomentosa* were separated into 4 groups. They were irrigated thoroughly and then withheld watering for 0 d, 3 d, 6 d, and 12 d separately. Leaves were collected on day 12. For dehydration, plants were moved out of the soil and dehydrated for 0 h, 1 h, 2 h, 3 h, 4 h, and 5 h. Leaves were collected after that. For salt treatment, twelve comparable pots were selected and gently pulled out and rinsed carefully. The plantlets were separated into four groups and cultured in 1/5 Hoagland nutrient solution for one week and then transferred into 1/5 Hoagland nutrient solution containing 200 mM NaCl for 0 h, 6 h, 12 h, and 24 h. The roots of each group were cut down after that. All these collected samples were frozen in liquid nitrogen immediately after being detached from plants and stored in an ultra-low temperature refrigerator (-80 °C) for RNA extraction.

### 2.6 RNA isolation and qRT-PCR analysis

Samples stored at -80 °C were ground into fine powders in liquid nitrogen. High-quality total RNA was isolated using the cetyltrimethylammonium bromide (CTAB) method according to the procedure reported previously (White et al., 2008). The quality of RNA (evaluating 28S and 18S ribosomal bands) was visualized by a 1% agarose gel and the quantity was measured by a NanoDrop One Microvolume UV-Vis Spectrophotometer (Thermo, West Palm Beach, FL). First-strand cDNA was synthesized using a HiScript II 1st Strand cDNA Synthesis Kit (Vazyme Nanjing, China), and the quantitative real-time PCR was performed using ChamQ SYBR qPCR Master Mix of Vazyme according to the manufacturer’s guidelines. The qRT-PCR assay was performed using a Quantagene q225 real-time PCR system (Kubo Technology, Beijing, China). Each sample contained three duplicates. The *R. tomentosa* Actin (*RtActin*) gene was used as an endogenous reference (He et al., 2018; Kanwar et al., 2023). Relative expression level of genes was determined by the 2^-ΔΔCT^ method (Livak and Schmittgen, 2001).

### 2.7 Generation of transgenic *Arabidopsis* and heat stress tolerance assay

The promoter of *AtUBQ10* was used to drive the coding sequence of *RtHSFA2a* and *RtHSFA2b* and the cassettes were inserted into a modified pCAMBIA1300 binary vector. *Agrobacterium tumefaciens* strain GV3101 carrying these overexpression vectors was used to transform *A. thaliana* (Col-0) by the floral dip method (Clough and Bent, 1998). T3 homozygous lines were used for further experiments.

For *Arabidopsis* heat stress treatment, 2-day-old germinated seedlings on 1/2 MS medium were transferred to a new plate and maintained at 22 °C for 3 days. They were suddenly exposed to 45 °C for 2.5 h without previous acclimation and recovered at 22 °C for another 3 days. Survival rate, biomass, and total chlorophyll content were analyzed to quantify the performances of different lines thereafter. For biomass and total chlorophyll content measurement, three shoots of plants were harvested and pooled as one sample. Chlorophyll was extracted with 95% ethanol till the plant turned white, and the absorbances of 665 nm and 649 nm were measured by a BioTek Epoch Microplate Spectrophotometer (BioTek, Santa Clara, CA). Total chlorophyll content was calculated as follows: Total chlorophyll (μg/g FW) = (18.16 × A_649_ + 6.65 × A_665_) × V ÷ FW [V: Extract volume (ml); FW: Fresh weight (g)].

### 2.8 Transient expression of *RtHSFA2a* and *RtHSFA2b* in *R. tomentosa*

Seedings of *R. tomentosa* growing in soil for 1 month were used for transient expression. *A. tumefaciens* (strain GV3101) carrying the overexpression vectors of *RtHSFA2a* and *RtHSFA2b* or empty vector were cultured in YEB medium overnight at 28 °C (200 rpm); they were centrifuged at 3500 rpm for 15 min and re-suspended by 1/2 MS medium (pH 5.8) supplied with 30 g/L sucrose, 200 μM acetosyringone and 0.01% silwet-77 to a final OD_600_ ∼ 0.8. *R. tomentosa* seedlings were gently removed from the soil and submerged in the infiltration medium. They were then vacuumized for 10 min and then re-planted in natural sands supplied with saturated water and covered with plastic wrap to keep humidity. They were kept in darkness for 24 hours and transferred into light for recovery. Three days after infiltration, samples were collected and used for RNA extraction and qRT-PCR analysis.

### 2.9 Promoter activity and transactivation assays

For the promoter activity assay of *RtHSFA2a* and *RtHSFA2b*, the promoter regions of *RtHSFA2a* and *RtHSFA2b* (2 kb upstream of the open reading frame) were cloned to drive the luciferase in the vector pGreenII-0800-LUC separately. CaMV35S::Ren was used as an internal control. These reconstructed vectors were transformed into *A. tumefaciens* strain GV3101 (pSoup) and used for *Nicotiana benthamiana* infiltration. Infected tobacco plants were kept in darkness for 48 hours. After that, some tobacco plants were treated by elevated temperature (37 °C) or dehydration for 2 h, others were kept at 25°C. Firefly luciferase was detected after that by a Nightshade evo LB985 In Vivo Plant Imaging System (Berthold Technologies, Bad Wildbad, Germany). Relative luciferase activity was quantified by the Dual-Luciferase Reporter Assay Kit of Vazyme.

For the transactivation assay, the promoters (2 kb upstream of the open reading frame) of putative targets of *RtHSFA2a* and *RtHSFA2b* were inserted into pGreenII-0800-LUC to drive luciferase. They were transformed into *A. tumefaciens* strain GV3101 (pSoup) and used as reporters. The overexpression vectors of *RtHSFA2a* and *RtHSFA2b* along with the empty vector were used as effectors and transformed into *A. tumefaciens* strain GV3101. These *A. tumefaciens* containing effectors and reporters were cultured in YEB medium overnight and used for *N. benthamiana* infiltration. Firefly luciferase and relative luciferase activity were detected as above.

### 2.10 Statistical analysis

All values were averaged based on three replicates in our study and are shown as mean ± s.d. Data analysis was performed using R (version 4.3.2, https://www.r-project.org/, last accessed on 2 March 2024). A one-way ANOVA analysis with Tukey’s HSD test was conducted for multiple comparisons (*p* < 0.05). Statistical differences were denoted by different lowercase letters.

## 3 Results

### 3.1 Identification of *RtHSF* genes in *R. tomentosa*

According to the identification of HMM and verification of SMART and HEATSTER, we finally confirmed that the *R. tomentosa* genome encodes 25 *RtHSF* members, and they were named according to the *AtHSF* gene names (Table 1). The 25 *RtHSF* genes could be categorized into three classes: HSFA, HSFB, and HSFC, each class contains 15, 8, and 2 members, respectively. Among these proteins, HSFB genes had the least amino acids (302 aa on average), while the HSFA genes had around 453 aa on average. The molecular weight ranged from 27.946 kDa (*RtHSFB5*) to 60.183 kDa (*RtHSFA9b*), and the pI ranged from 4.55 (*RtHSFA8*) to 8.73 (*RtHSFB3*) (Table 1). These proteins were all predicted localized in the nucleus (Table 1).

**Table 1.** Summary information of 25 *RtHSF* genes of *R. tomentosa*. P.I, iso-electric point; MW, molecular weight of amino acid sequence.

| Gene ID | Gene Name | Protein length | P.I | MW | Predicted Subcellular Locations |
| --- | --- | --- | --- | --- | --- |
| Rt01g30040.1 | *RtHSFA3* | 508 | 5.63 | 56752.93 | Nucleus |
| Rt01g39160.1 | *RtHSFA4c* | 436 | 5.2 | 49438.15 | Nucleus |
| Rt01g44380.1 | *RtHSFB3* | 263 | 8.43 | 29732.34 | Nucleus |
| Rt01g50380.1 | *RtHSFB2b* | 319 | 4.82 | 34602.75 | Nucleus |
| Rt01g52500.1 | *RtHSFA1c* | 512 | 5.07 | 56593.5 | Nucleus |
| Rt02g08590.1 | *RtHSFA8* | 397 | 4.55 | 44983.71 | Nucleus |
| Rt03g05150.1 | *RtHSFC1b* | 362 | 7.74 | 39810.66 | Nucleus |
| Rt03g06670.1 | *RtHSFA9a* | 474 | 5.16 | 52661.39 | Nucleus |
| Rt03g06680.1 | *RtHSFA9b* | 537 | 5.78 | 60183.09 | Nucleus |
| Rt03g10240.1 | *RtHSFA2b* | 369 | 4.73 | 41732.68 | Nucleus |
| Rt03g44450.1 | *RtHSFC1a* | 317 | 5.32 | 34988.35 | Nucleus |
| Rt03g47450.1 | *RtHSFA9c* | 463 | 5.02 | 51004.91 | Nucleus |
| Rt03g50460.1 | *RtHSFA2a* | 397 | 5.73 | 44435.81 | Nucleus |
| Rt04g06900.1 | *RtHSFB5* | 243 | 7.72 | 27946.18 | Nucleus |
| Rt04g41480.1 | *RtHSFA1a* | 506 | 5.54 | 56268.25 | Nucleus |
| Rt05g00320.1 | *RtHSFA4a* | 423 | 5.01 | 48067.24 | Nucleus |
| Rt05g44170.1 | *RtHSFB2c* | 338 | 4.78 | 36312.39 | Nucleus |
| Rt05g49220.1 | *RtHSFA1b* | 514 | 4.77 | 56380.62 | Nucleus |
| Rt06g33750.1 | *RtHSFB4a* | 367 | 8.44 | 40743.12 | Nucleus |
| Rt08g25280.1 | *RtHSFA6a* | 406 | 5.04 | 45807.31 | Nucleus |
| Rt08g37810.1 | *RtHSFA5* | 507 | 5.82 | 56885.25 | Nucleus |
| Rt09g22770.1 | *RtHSFB1* | 294 | 6.98 | 31886.33 | Nucleus |
| Rt10g41750.1 | *RtHSFB4b* | 304 | 6.76 | 34735.1 | Nucleus |
| Rt11g02760.1 | *RtHSFB2a* | 288 | 8.73 | 32103.25 | Nucleus |
| Rt11g55470.1 | *RtHSFA6b* | 354 | 5.38 | 40071.31 | Nucleus |

### 3.2 Chromosomal spread and phylogenetic analysis of *RtHSF* members

All these *RtHSF* genes were successfully mapped to the linkage groups of the *R. tomentosa* genome, however, they were distributed unevenly on the chromosomes (Figure 1). Chromosomes 1 and 3 encoded the largest number of *RtHSF* genes, 5 and 6 members respectively, followed by chromosome 5 (3 *RtHSF* genes), while chromosome 7 had no *RtHSF* member. Over half of the *RtHSF* genes were located at the ends of chromosomes.

**Figure 1.**
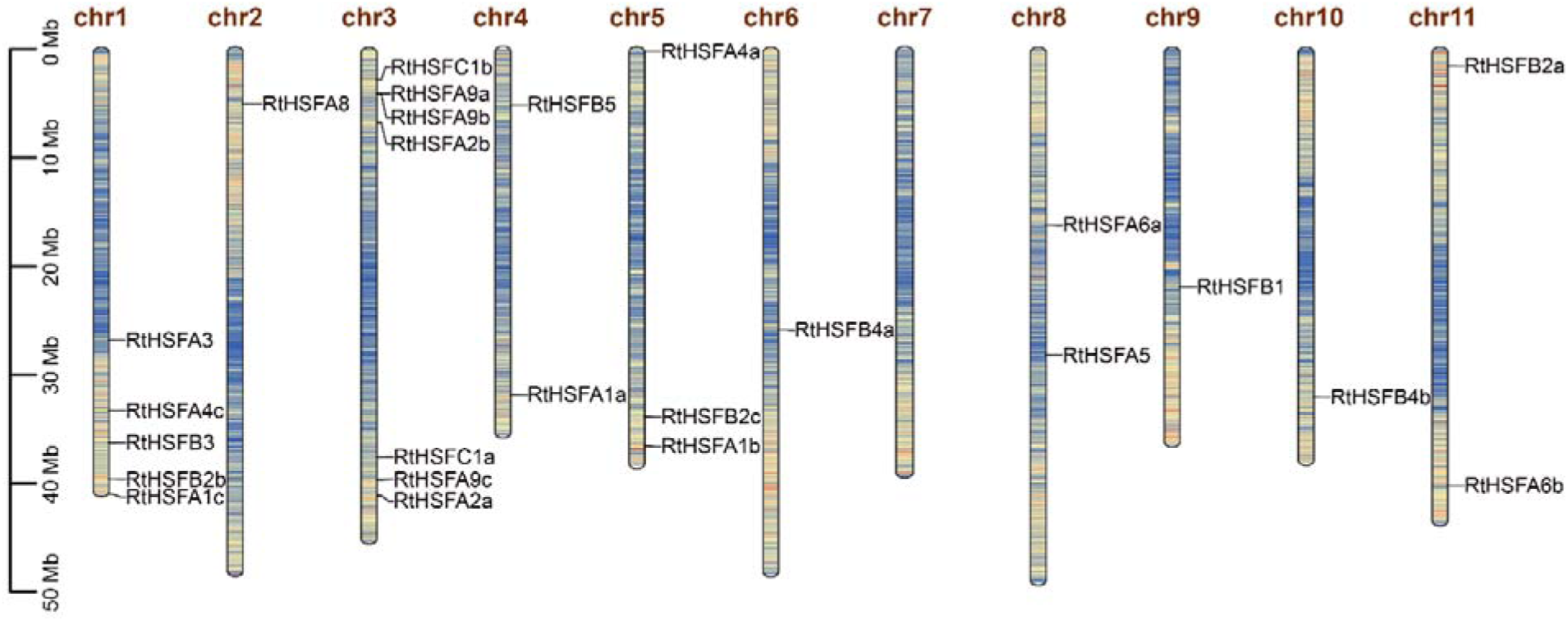
Chromosomal locations of *RtHSF* family members. The length and gene density of 11 chromosomes were presented by ribbons. Blue lines represented a relatively low gene density.

In order to study the evolutionary relationships of the HSF gene family of *R. tomentosa*, we constructed a phylogenetic tree using the Maximum likelihood (ML) approach together with the HSF genes of *A. thaliana* and *E. grandis*. Based on the bootstrap values and the evolutionary link with *A. thaliana* and *E. grandis*, HSFA could be further divided into 9 subclasses (HSFA1-A9), HSFB could be divided into 5 subclasses (HSFB1-B5), and HSFC contained only 1 subclass (HSFC1) (Figure 2). We observed that *RtHSF* and *EgHSF* families contained no HSFA7 members. Remarkably, the *RtHSF* gene family had 3 members in subclass HSFA9 and HSFB2, and two members in subclass HSFB4, while *AtHSFs* contained 1, 2, and 1 member in these subclasses separately. We also identified one member in *HSFB5* both in the *R.tomemtosa* and *E. grandis* genome, whereas it was absent in the *A. thaliana* genome.

**Figure 2.**
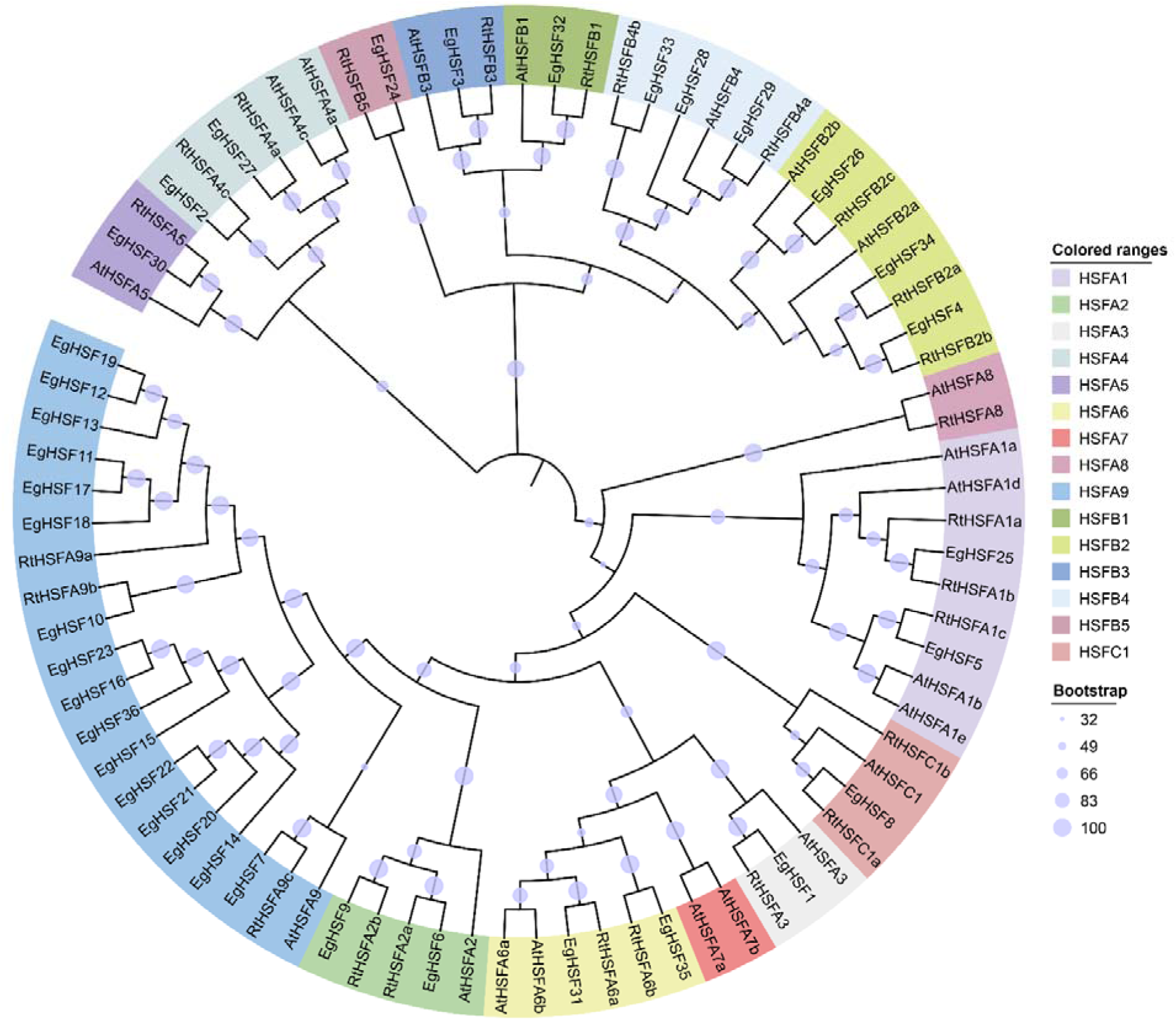
Phylogenetic relationship of HSF family of *R. tomentosa*, *A. thaliana*, and *E. grandis*. Amino acid sequences of these genes were used for tree construction. These genes could be classified into 9 clades and were noted by different colors. The size of the circle on the branch represented the Bootstrap value based on 1000 iterations.

### 3.3 Synteny and gene duplication analyses of *RtHSF* genes

Gene duplication or whole-genome duplication as an important raw genetic material provides functional novelty in animals, fungi, and other organisms, especially in plant evolution and distribution (Jiao et al., 2011). We identified 9 pairs of syntenic genes in the *RtHSF* family as the red lines indicated (Figure 3a). Noteworthy, *RtHSFA2a* had synteny with *RtHSFA9a* and *RtHSFA9c*, and *RtHSFA2*b had synteny with *RtHSFA9a*. We further analyzed the force that drove the expansion of the *RtHSF* gene family using DupGen_finder. Results showed that all 9 pairs of these paralogous genes were the results of whole-genome duplication (WGD) events (Table 2). To explore the evolutionary constraints of the *RtHSF* genes, we calculated the Ka (nonsynonymous substitution), Ks (synonymous substitution), and Ka/Ks ratios of the duplicated genes. All these gene pairs had a Ka/Ks ratio of less than 1 (Table 2), meaning that the *RtHSF* gene family has undergone strong purifying selection pressure in the course of evolution.

**Figure 3.**
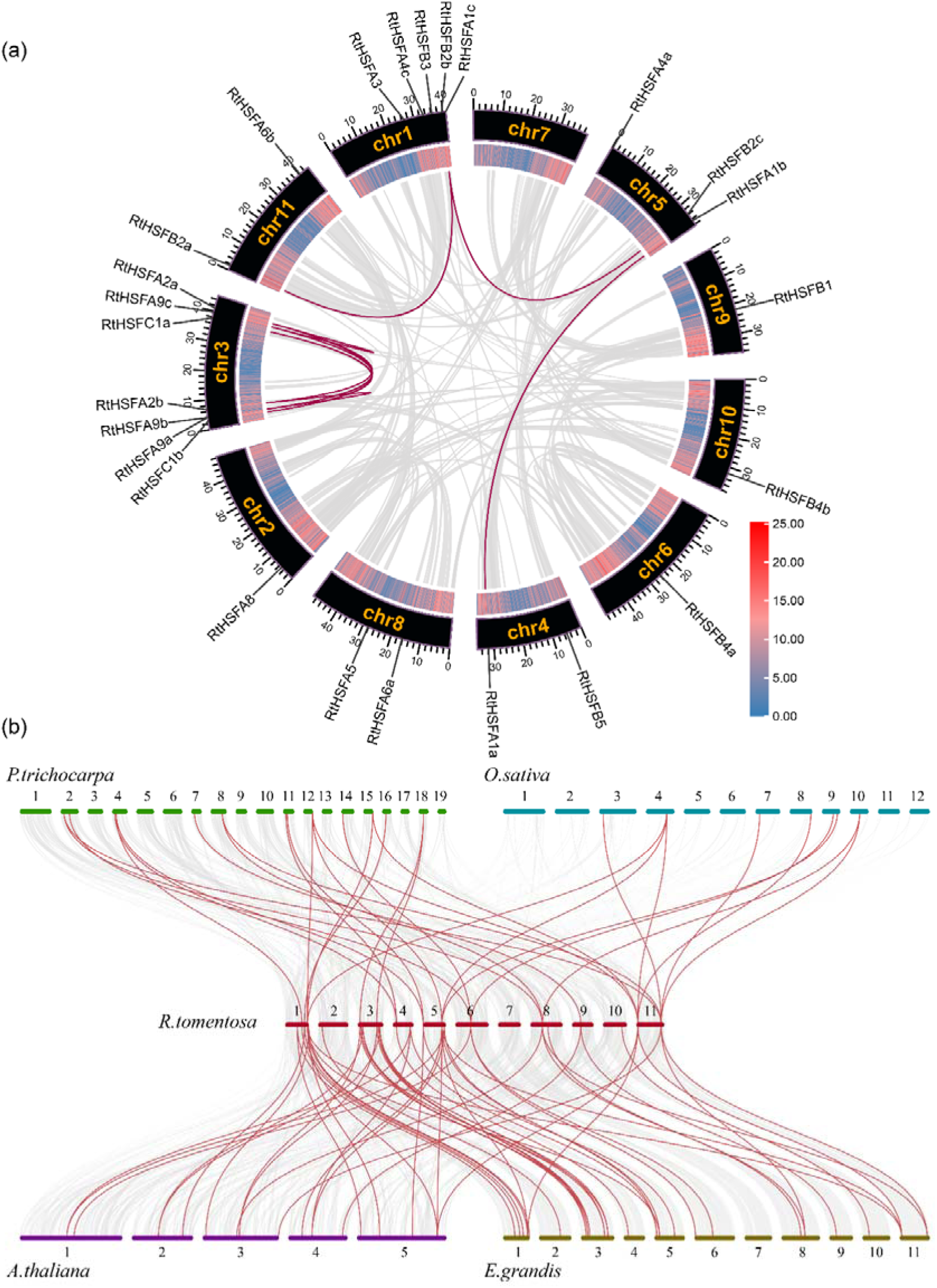
Synteny analysis of *RtHSF* family members. (a) Synteny analysis in *R. tomentosa*. Red lines indicated the syntenic gene pairs of *RtHSFs*. Legend indicated the gene intensity on chromosomes of *R. tomentosa*. (b) Synteny analysis of *R. tomentosa* with *P. trichocarpa*, *O. sativa*, *A. thaliana*, and *E. grandis*. Grey lines indicated the synteny blocks between paired genomes and HSF genes were highlighted by red lines.

**Table 2.** Gene duplication type and selection pressure analyses of the orthologous *RtHSF* genes. WGD: whole-genome duplication.

| Gene pairs | Ka | Ks | Ka/Ks | Selective type | Gene duplication type |
| --- | --- | --- | --- | --- | --- |
| RtHSFB2a-RtHSFB2b | 0.85562802 | 0.891245691 | 0.96003608 | Purifying | WGD |
| RtHSFB2b-RtHSFB2c | 0.981897897 | 1.020038597 | 0.962608571 | Purifying | WGD |
| RtHSFC1a-RtHSFC1b | 0.965769936 | 1.334868089 | 0.723494662 | Purifying | WGD |
| RtHSFA9a-RtHSFA9c | 0.503266788 | 2.118497221 | 0.23755839 | Purifying | WGD |
| RtHSFA2a-RtHSFA2b | 0.292002401 | 1.147828104 | 0.254395584 | Purifying | WGD |
| RtHSFA9a-RtHSFA2b | 0.538857001 | 1.518311843 | 0.354905353 | Purifying | WGD |
| RtHSFA9c-RtHSFA2a | 0.537838446 | 1.675699387 | 0.320963563 | Purifying | WGD |
| RtHSFA9a-RtHSFA2a | 0.634209319 | 1.535172277 | 0.413119315 | Purifying | WGD |
| RtHSFA1a-RtHSFA1b | 0.249292949 | 1.548522757 | 0.160987591 | Purifying | WGD |

To obtain a deeper understanding of the HSF gene family phylogeny, we constructed a synteny analysis of the HSF gene family across species including *R. tomentosa*, *A. thaliana*, *O. sativa*, *P. trichocarpa*, and *E. grandis*. Result indicated that *RtHSFs* were distantly related to *O.sativa*, while had close origin relationships with *A. thaliana*, *P. trichocarpa*, and especially *E. grandis*, a near relative of *R. tomentosa* (Figure 3b).

### 3.4 Sequence structure and pattern of conserved motifs

The number of exons varied from 2-4 in *RtHSF* genes, and most of them (21 members) contained two exons and one introns (Figure 4b). *RtHSFA6a*, *RtHSFA9b*, and *RtHSFA3* contained three exons and two introns, whilst *RtHSFA1a* possessed the largest number of exons and introns, with four exons and three introns. The longest gene length was observed in *RtHSFA5* for it had a large intron, followed by *RtHSFA1b*. We observed that the number and arrangement of exons did not appear to be closely related to the evolutionary relationships except for a few clades. Some affinitive genes in evolution had more similar lengths of introns and exons (*RtHSFA4a* with *RtHSFA4b*, and *RtHSFC1a* with *RtHSFC1b)*, while other clades exhibited considerable variations in gene structure (Figure 4a-b). For example, *RtHSFA1a* contained four exons, while *RtHSFA1b* and *RtHSFA1c* contained two. We then submitted the amino acid sequences to the MEME tool to identify the conserved motifs of RtHSF genes. Ten motifs were predicted in these RtHSF genes (Figure 4c). Motifs 1-3 were identified as distinctive structural motifs of HSF and presented in all RtHSF genes except RtHSFA9a (lacking motif 2) and RtHSFB2b (lacking motif 3). Motif 4 existed in all the HSFA class genes as well as RtHSFC1b, and motif 5 was found in all the HSFA class and RtHSFC1a. Motif 7 was specific to the HSFA class, and motif 6 was specific to the HSFB class. Affinitive genes contained similar genetic composition and arrangement (Figure 4a, c), demonstrating the highly conservative relationships during their evolution at the protein level.

**Figure 4.**
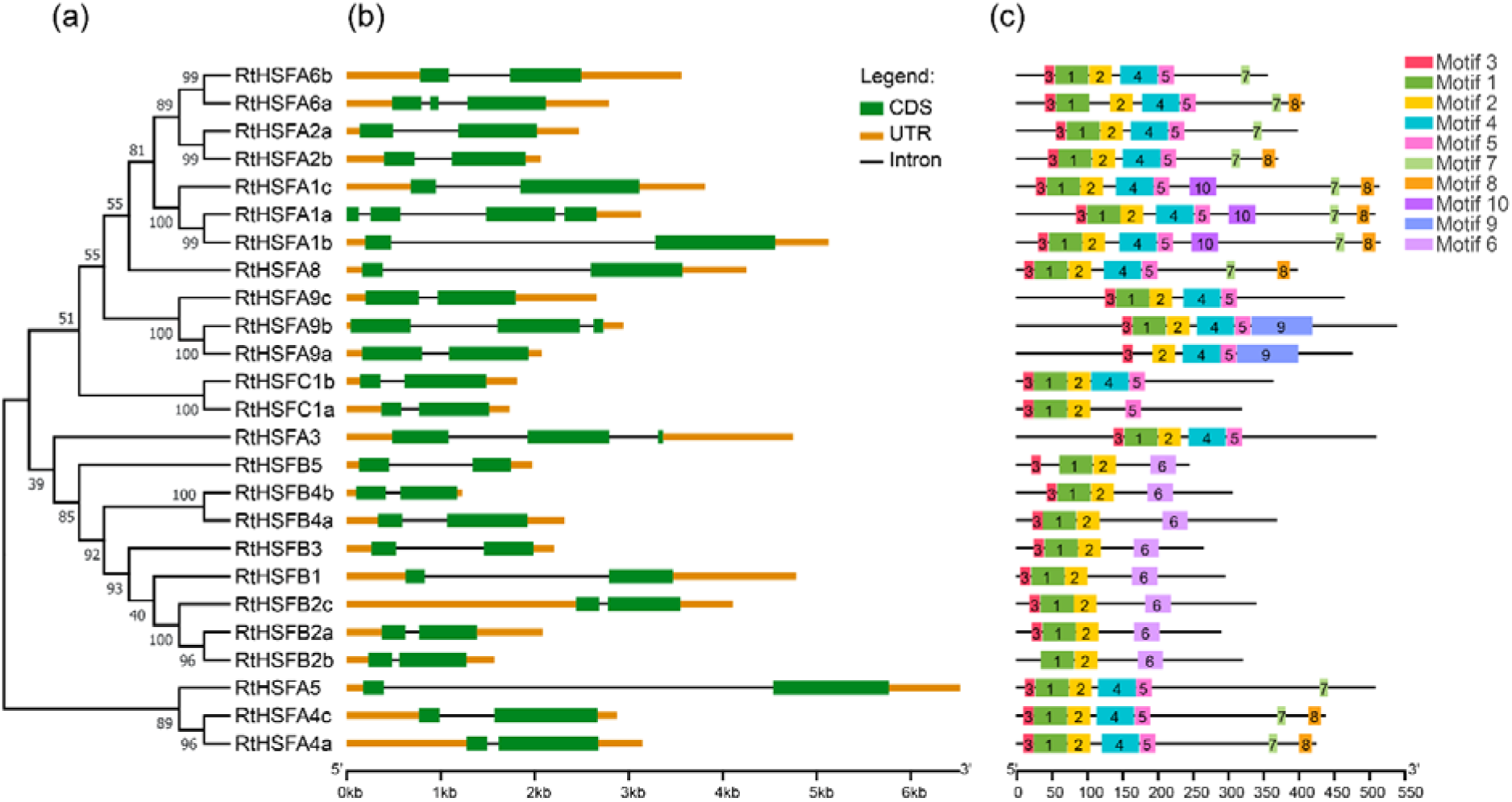
Structures of *RtHSF* genes. (a) Evolutionary relationships of RtHSFs. Amino acid sequences of RtHSF genes were used to build the phylogenetic tree and ML method with 1000-boostrap replications was applied. The number on each branch indicated the bootstrap value. (b) Gene structure of *RtHSFs*. The exon, intron, and UTR (Untranslated Regions) of *RtHSFs* were displayed by GSDS. (c) Distribution of conserved motifs throughout the RtHSFs. Amino acid sequences of RtHSFs were subjected to the analysis of the MEME suite and ten motifs were predicted.

### 3.5 *Cis-acting* element analysis of the promoter region of *RtHSFs*

Gene functions in organisms largely depend on their expression pattern. The *cis-acting* elements in the promoter regions, which are the specialized binding sites of proteins, are responsible for regulating transcription. We analyzed the promoter regions of these 25 *RtHSF* genes (2 kb upstream of the open reading frame), and 20 *cis-acting* elements relevant to the expression regulation of *RtHSFs* were retrieved. Results showed that *RtHSFs* might take roles during the whole lifecycle of *R. tomentosa* (Figure 5a). Roughly, these *cis-acting* elements could be categorized into growth and development-related elements, phytohormone-responsive elements, and stress-responsive elements. The growth and development-related elements include endosperm expression regulatory element, meristem expression regulatory element, circadian control related elements, etc. The stress-responsive elements contained anaerobic response elements, oxidative response elements (as-1), low-temperature response elements, wound responsive element, defense and stress-responsive element, etc. There were also a great lot of elements related to major phytohormones including ABA, ethylene, auxin, gibberellin, salicylic acid, and MeJA (Figure 5a-b). *RtHSFs* of varying types shared the presence of the light-responsive element, abscisic acid-responsive element; they also possessed MYB and MYC recognition sites, two critical gene families related to growth and development and stress response (Figure 5b). Noteworthy, there existed a large number of ABA-responsive elements in *RtHSFA2a* (15), *RtHSFA4c* (18), *RtHSFA9a* (13), *RtHSFA9b* (12), *RtHSFB2c* (18), and *RtHSFB4b* (24), compared with their homologous genes (Figure 5b). Same with the ABA-responsive element, some *RtHSFs* were specialized in response to MeJA. For example, *RtHSFA1b* had 16 MeJA responsive elements, while the *RtHSFA1a* and *RtHSFA1c* had 8 and 4 respectively; *RtHSFA9c* had 12 MeJA responsive elements, whilst *RtHSFA9a* and *RtHSFA9c* possessed 0 and 2, separately (Figure 5b).

**Figure 5.**
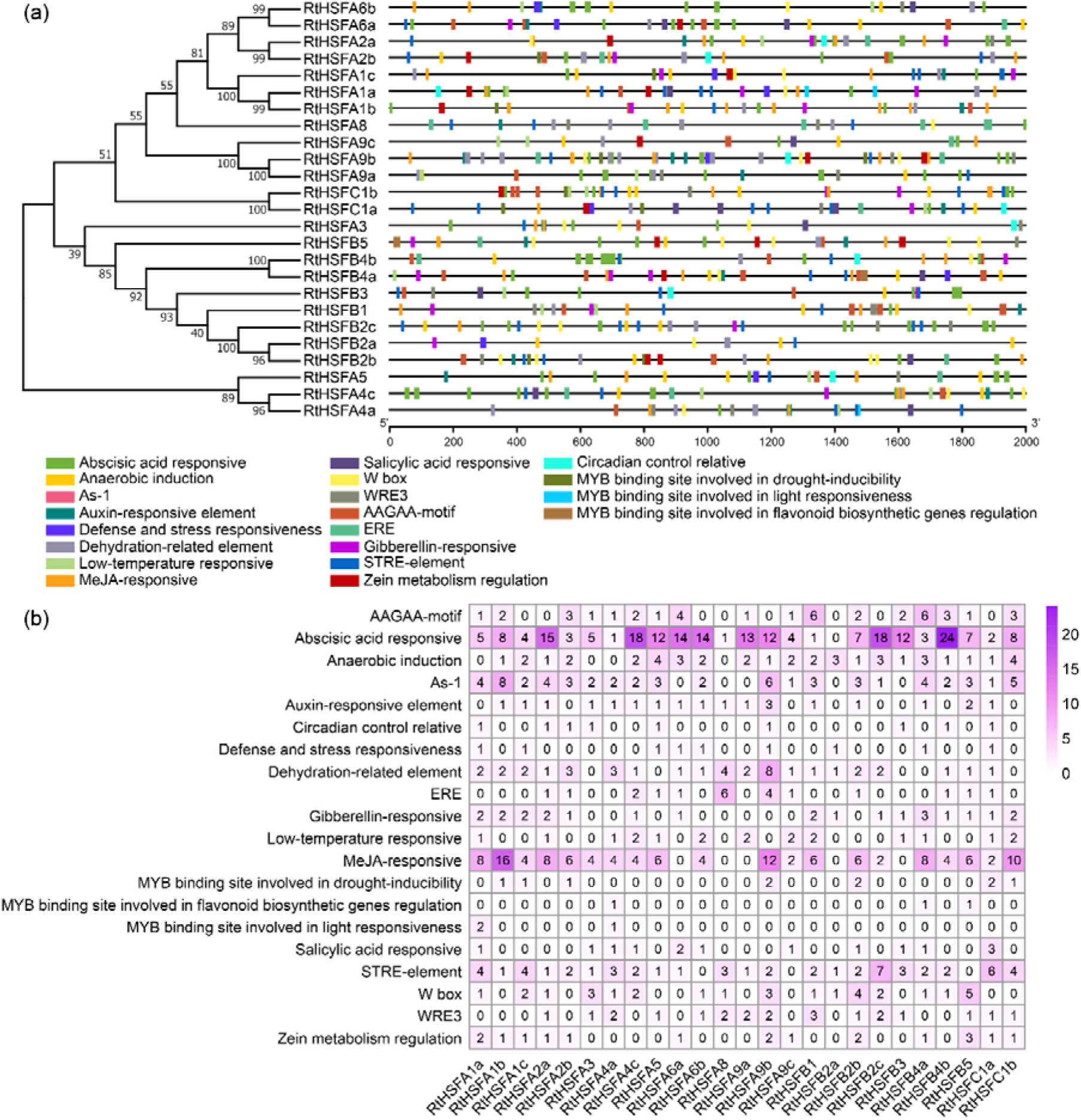
The *cis-acting* elements analysis of the promoter of *RtHSF* genes. The promoter sequences (2 kb upstream of the open reading frame) of 25 *RtHSF* genes were subjected to the analysis of PlantCARE and 25 types of *cis*-*acting* elements were retrieved. (a) Annotation and distribution of *cis-acting* elements in the promoter regions of *RtHSF* genes. (b) Statistical analysis of *cis-acting* elements of *R. tomentosa* HSF gene promoters. The number of each grid denoted the total number of *cis-acting* elements in the promoter of the relevant gene.

### 3.6 Expression profile of *RtHSF* genes in different tissues

RNA-seq-based transcriptome data of *R. tomentosa* were explored to reveal the expression pattern across various organs including root, stem, leaf, flower, and different maturing statuses of fruit (green fruit, yellow fruit, and red fruit) (Yang et al., 2024). All of the *RtHSFs* could be detected in the sequenced tissues except for HSFB4b, which was absent in the flower (Figure 6). They exhibited differential expression patterns among vegetative and reproductive tissues, which could be divided into 6 groups. *RtHSFA1b*, *RtHSFA5*, *RtHSFA4c*, *RtHSFB1*, *RtHSFB3*, *RtHSFB5*, and *RtHSFC1a* showed higher expression levels in root. *RtHSFA2b*, *RtHSFA4a*, *RtHSFB2b*, *RtHSFB2c,* and *RtHSFC1b* showed higher expression levels in flower. *RtHSFA6b* and *RtHSFB2a* had a higher mRNA abundance in green fruit (Figure 6). Subclass *RtHSFA9* demonstrated high expression in fruits and low expression in flower and vegetative tissues. Detailedly, *RtHSFA9a* had a higher expression level in green fruit, while *RtHSFA9b* was elevated in yellow fruit and red fruit (Figure 6).

**Figure 6.**
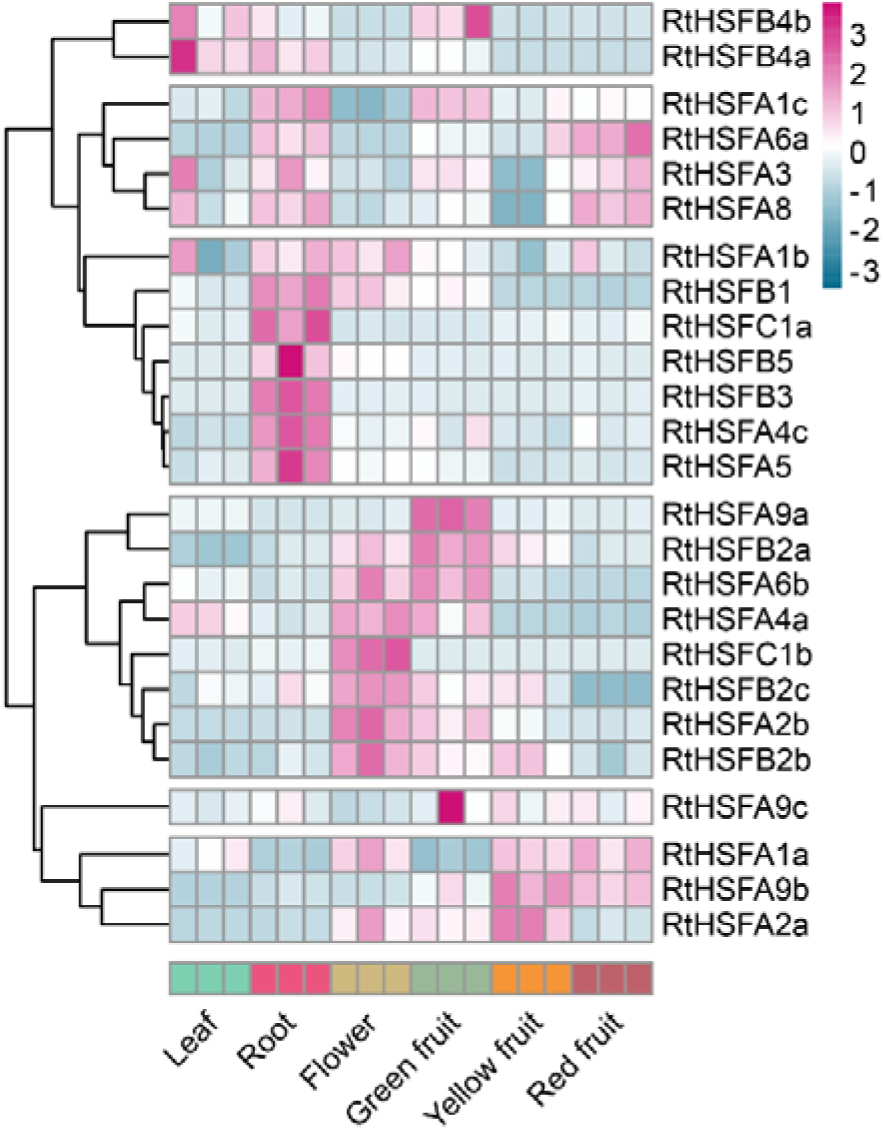
Heat map of the tissue-specific expression profile of *RtHSFs*. Root, stem, leaf, flower, green fruit, yellow fruit, and red fruit were collected and used for RNA-Seq previously (Yang et al., 2024). Each sample contained three duplications. Transcripts per million values (TPM) were used for heat map plotting.

### 3.7 Expression pattern of *RtHSF* genes under heat stress conditions

We selected *RtHSF* genes from four subclasses to examine their expression levels under temporal sudden heat stress. These genes were the results of gene duplication events, and we wonder whether there existed divergences in gene functions under heat stress conditions. As shown in Figure 7, both of the two *RtHSFA2* were strongly induced by detrimental elevated temperature and reached the highest relative mRNA abundance after 1 h treatment. However, *RtHSFA2b* exhibited a prodigious response to heat stress, reaching a peak of around 2923-fold at 1 h (Figure 7a), whereas the relative mRNA abundance of *RtHSFA2a* was 60-fold of the control at the same time (Figure 7b). The mRNA abundances of *RtHSFA2a* and *RtHSFA2b* descended 6 h after heat treatment, however, they still maintained relatively high levels (around 51-and 1689-fold of the control). Similar to *RtHSFA2*, all members of the *RtHSFA9* subclass exhibited different expression patterns under heat stress. The expression of *RtHSFA9a* was not affected by heat stress (Figure 7c), while *RtHSFA9b* and *RtHSFA9c* slightly and gradually responded to high temperature, reaching peaks at 6 h (around 1.6-and 2.9-fold upregulation) (Figure 7d-e). All of the *RtHSFB2* members, as well as *RtHSFC1b*, were induced by heat stress gradually, reaching around 3∼8-fold at 6 h (Figure 7f-i).

**Figure 7.**
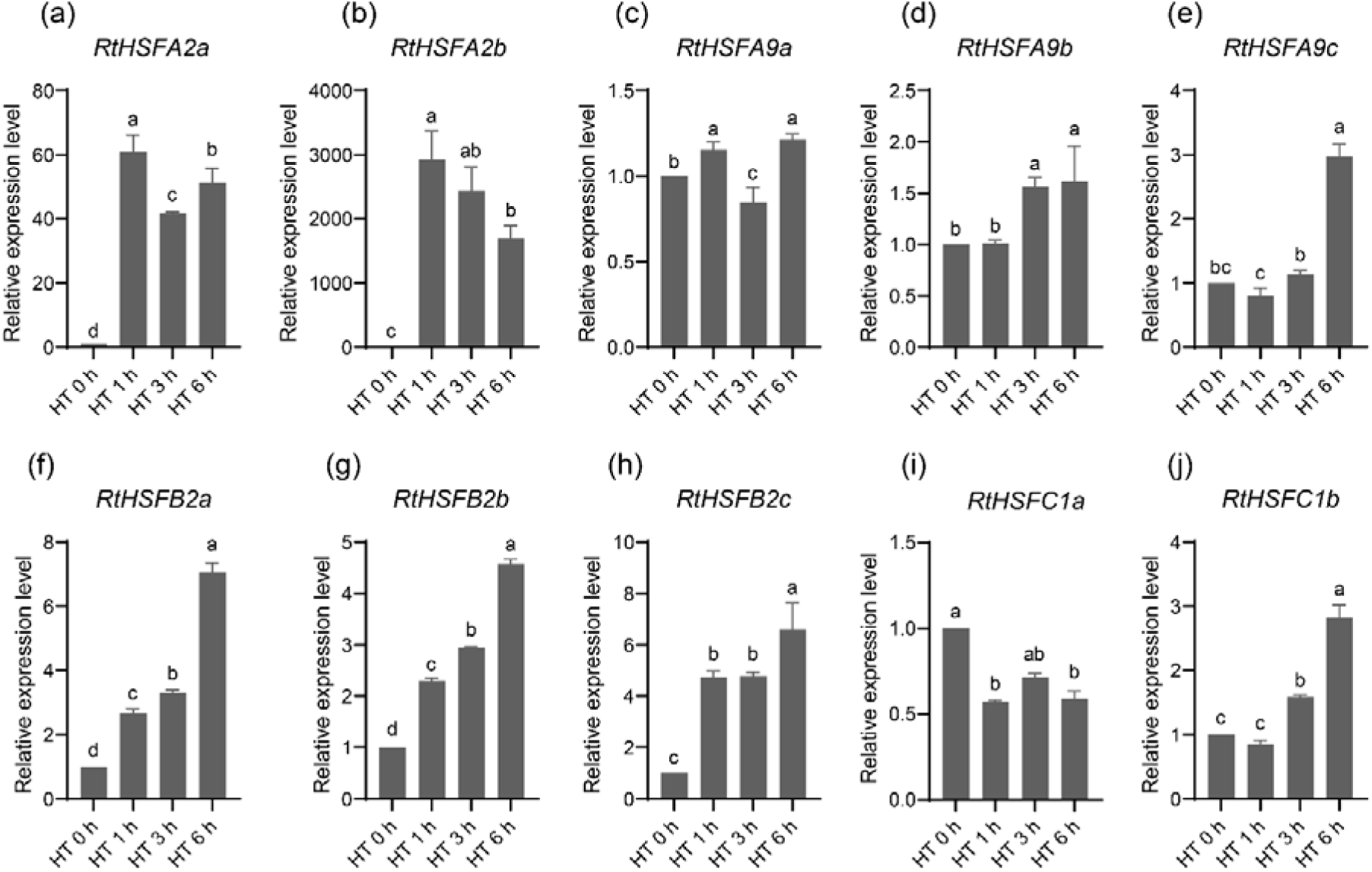
Expression patterns of *RtHSF* genes in response to heat stress. Representative *RtHSF* genes of subclasses with gene duplications were selected for quantification to detect heat stress responsive patterns. Leaves after being subjected to 45 °C for 0 h, 1 h, 3 h, and 6 h were collected for qRT-PCR. The sample of 0 h was used as a control. Values were mean ± SD (n = 3). One-way ANOVA followed by post-hoc Tukey’s HSD was applied to evaluate the significance among different heat treatment durations (*p* < 0.05). Samples sharing the same letters showed no significant difference.

### 3.8 Expression pattern of *RtHSFA2a* and *RtHSFA2b* in response to abiotic stresses

The amino acid sequences of *RtHSFA2a* and *RtHSFA2b* showed a large part variation, sharing only 51% identity (Figure 8a). Considering the promoter of *RtHSFA2a* contained 15 AREB *cis-acting* elements and only 3 in that of *RtHSFA2b* (Figure 5), we speculated that *RtHSFA2a* and *RtHSFA2b* might exist functional divergences during evolution to cope with different stresses specifically. We thus analyzed the expression profile of *RtHSFA2a* and *RtHSFA2b* under various abiotic stresses. Different from heat stress, *RtHSFA2a* was more sensitive to drought stress (Figure 8b). Although both the mRNA abundance of *RtHSFA2a* and *RtHSFA2b* increased gradually as the drought stress aggravated and reached a peak after 12 days’ treatment, the relative expression level of *RtHSFA2a* was much higher than that of *RtHSFA2b* (22-fold versus 2.6-fold). The same tendency appeared in dehydration treatment (Figure 8c). Similar to drought stress, *RtHSFA2a* and *RtHSFA2b* both responded to salt stress gradually, whilst the mRNA abundance of *RtHSFA2a* reached a peak of 32.7-fold after 24 h when subjected to salt stress, and *RtHSFA2b* reached 18-fold at that point (Figure 8d). Moreover, *RtHSFA2b* showed a weak correlation with cold stress, while *RtHSFA2a* slightly responded to cold stress (Figure 8e).

**Figure 8.**
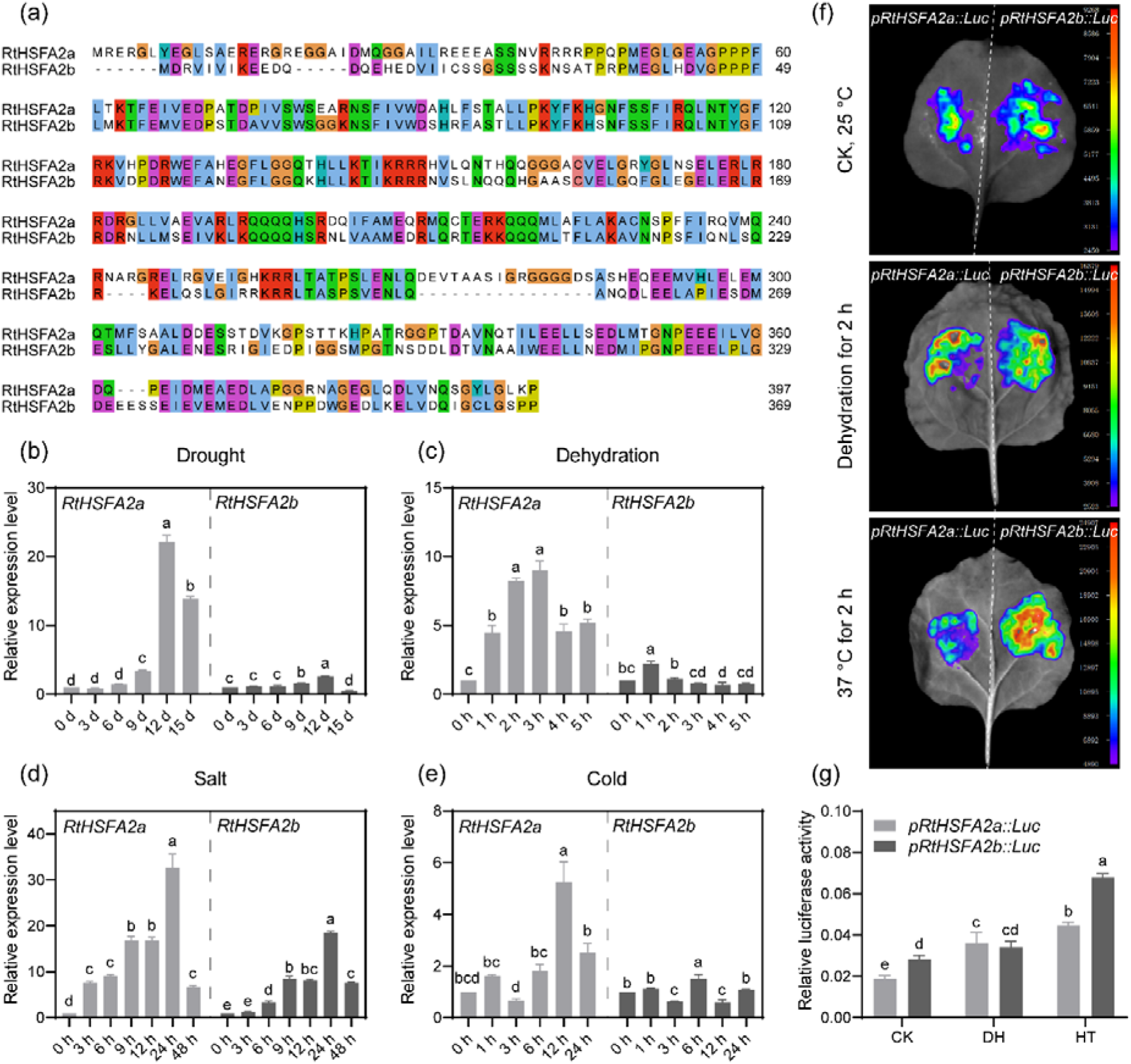
Comparison of amino acid sequence and expression patterns of *RtHSFA2a* and *RtHSFA2b* in response to abiotic stresses. (a) Amino acid sequence alignment of *RtHSFA2a* and *RtHSFA2b*. (b-e) Relative mRNA abundance of *RtHSFA2a* and *RtHSFA2b* under drought, dehydration, salt, and cold stresses. Samples of 0 h were used as controls. (f) Promoter activities of *RtHSFA2a* and *RtHSFA2b* under normal, dehydration, and heat stress conditions in tobacco leaves. (g) Statistical analysis of the relative Luc/Ren values in (f). CK, control; DH, dehydration for 2 h; HT, 37 °C for 2 h. Values were mean ± SD (n = 3). Statistical significance was calculated by one-way ANOVA with Tukey’s HSD test (*p* < 0.05). Significant differences were denoted by different letters.

We further compare the transcriptional activities of the *RtHSFA2a* and *RtHSFA2b* promoters under normal, dehydration, and heat stress conditions by luciferase assay. As the results shown in Figure 8f-g, the relative activity of the *RtHSFA2b* promoter was slightly higher than that of *RtHSFA2a* under normal conditions. When treated by dehydration, the promoter activity of *RtHSFA2a* increased and showed no significant difference compared with that of *RtHSFA2b*. After two hours of heat stress treatment (37 °C), the activity of *RtHSFA2a* and *RtHSFA2b* promoters were all elevated, while the *RtHSFA2b* promoter exhibited a significantly higher transcriptional activity than that of *RtHSFA2a* (Figure 8f-g).

### 3.9 Performance of *Arabidopsis* with overexpressed *RtHSFA2a* and *RtHSFA2b* under heat stress

As *RtHSFA2a* and *RtHSFA2b* varied in response to heat stress, we wonder whether there existed functional divergences in these two homologous genes under heat stress conditions. We then overexpressed them in *Arabidopsis* and evaluated their performance under acute heat stress. Results showed that although overexpressing *RtHSFA2a* conferred thermotolerance to *Arabidopsis*, plants with overexpressed *RtHSFA2b* (OEA2b line) performed significantly better than those with *RtHSFA2a* (OEA2a line) and Col-0 (Figure 9a). Plants of OEA2b lines showed no mortality after heat stress, while only 29-33% of OEA2a and 20% of Col-0 plants survived respectively (Figure 9c). In addition, OEA2b lines had higher biomass and total chlorophyll content after heat stress than those of OEA2a and Col-0 lines (Figure 9d).

**Figure 9.**
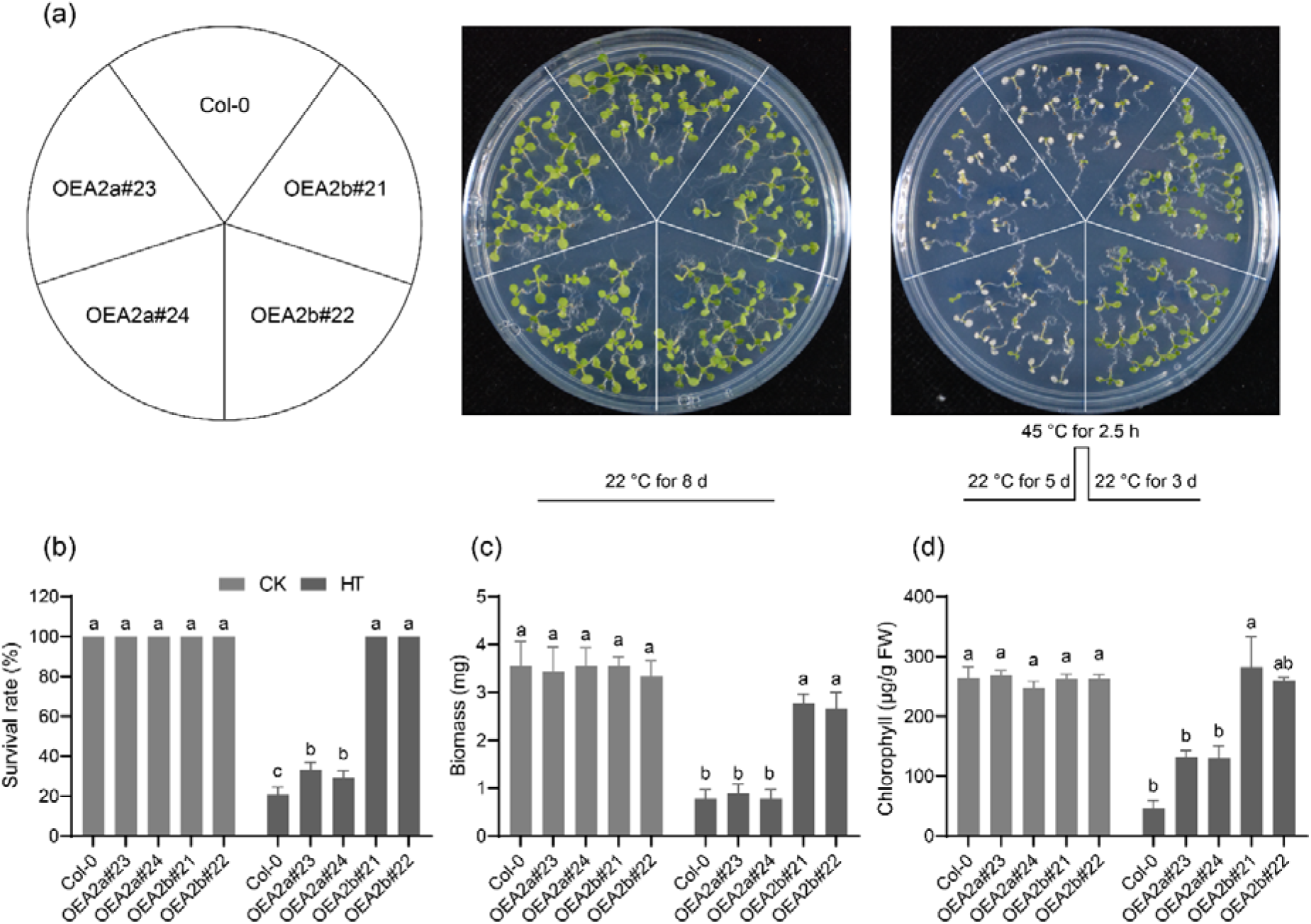
Performance of *Arabidopsis* with overexpressed *RtHSFA2a* and *RtHSFA2b* under acute heat stress. (a) Phenotypes of *Arabidopsis* of Col-0, OEA2a, and OEA2b lines under normal and heat stress conditions. Five-day-old plants were subjected to 45 °C for 2.5 h thereafter recovered at 22 °C for 3 d. Photographs of 8-day-old plants were taken. OEA2a: *Arabidopsis* plants with overexpressed *RtHSFA2a*. OEA2b: *Arabidopsis* plants with overexpressed *RtHSFA2b*. Col-0 was used as a control. (b-d) Survival rate, biomass, and total chlorophyll content of different *Arabidopsis* lines under normal and heat stress conditions. Values were mean ± SD (n = 3). One-way ANOVA followed by post-hoc Tukey’s HSD was applied to evaluate the significance among different heat treatment durations (*p* < 0.05). Samples sharing the same letters showed no significant difference.

### 3.10 *RtHSFA2a* and *RtHSFA2b* functioned differently in activating heat-responsive genes in *R. tomentosa*

Transient transformation is a simple, rapid, and effective method of vast potential in gene function investigation (Li et al., 2021; Zheng et al., 2021). We transiently overexpressed *RtHSFA2a* and *RtHSFA2b* in *R. tomentosa* seedlings, and the empty vector (VT) and WT were used as controls. We detected around 27-fold overexpression of *RtHSFA2a* in the OxHSFA2a line and around 20-fold overexpression of *RtHSFA2b* in the OxHSFA2b line (Figure 10a-b). We find that *RtHSFA2a* was slightly up-regulated in the OxHSFA2b line (Figure 10b), while *RtHSFA2b* was greatly transcriptional activated in the OxHSFA2a line, reaching around 10-fold of WT (Figure 10a). These indicated that *RtHSFA2a* might act as an activator of *RtHSFA2b*. We then quantified some potential target genes of *RtHSFA2a* and *RtHSFA2b*, including *RtHSP17.4*, *RtHSP21.3*, *RtHSP21.8*, *RtHSP26.5*, and *RtHSP70*. All these genes were upregulated in OxHSFA2a and OxHSFA2b lines compared with WT and VT controls (Figure 10c-g). We also measured another two putative targets of *RtHSFA2a* and *RtHSFA2b*: RtSKIP27, an F-box E3 ubiquitin ligase, and *RtZAT10*, a zinc-finger transcription factor known as an essential gene in ROS scavenging (Li et al., 2023). They were all up-regulated by over-accumulated *RtHSFA2a* and *RtHSFA2b*, however, differed from other genes, they had higher expression levels in the OxHSFA2b line, reaching around 2.4-fold and 3.1-fold versus 2.1-fold and 2.5-fold in the OxHSFA2a line respectively (Figure 10h-i).

**Figure 10.**
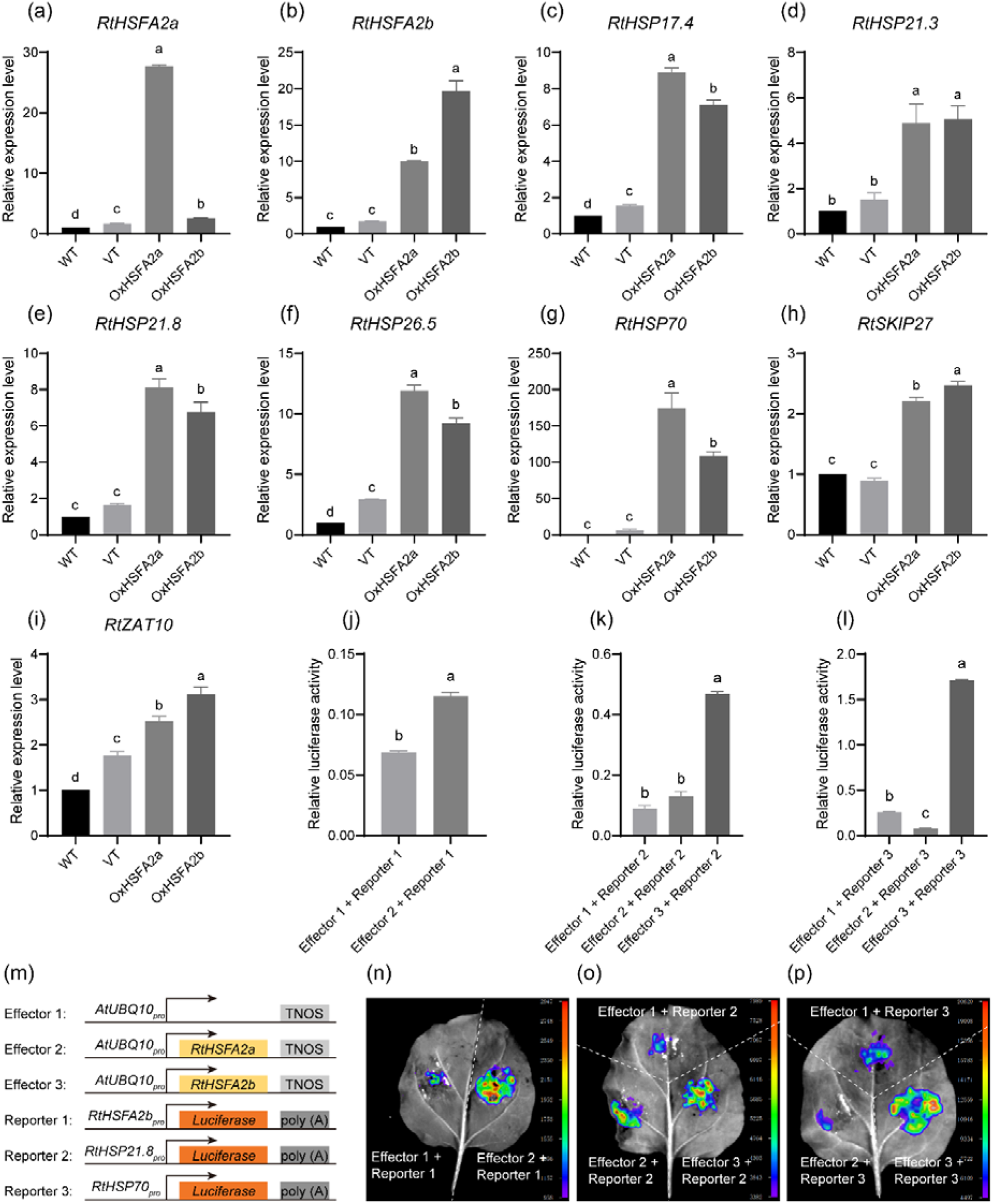
The regulatory network of *RtHSFA2a* and *RtHSFA2b* in *R. tomentosa*. *RtHSFA2a* and *RtHSFA2b* were transiently overexpressed in *R. tomentosa* seedlings and then samples were collected for qRT-PCR. WT, wild-type *R. tomentosa* seedlings without any treatment. VT, seedlings transformed with empty vector. OxHSFA2a, seedlings overexpressing *RtHSFA2a*. OxHSFFA2b, seedlings overexpressing *RtHSFA2b*. WT and VT were used as controls. (a-b) Quantifications of *RtHSFA2a* and *RtHSFA2b* expression levels in the transient expression lines of *R. tomentosa*. (c-i) Relative mRNA abundance of heat shock response genes in the transient expression lines of *R. tomentosa*. (j and n) Transactivation assay of the *RtHSFA2b* promoter by *RtHSFA2a* (n) and the relative luciferase activity (j). (k and o) Transactivation assay of the *RtHSFA2a* and *RtHSFA2b* to *RtHSP21.8* promoter (o) and the relative luciferase activity (k). (l and p) Transactivation assay of the *RtHSFA2a* and *RtHSFA2b* to *RtHSP70* promoter (p) and the relative luciferase activity (l). (m) Schematic of constructed plasmids used for the transactivation assays in *N. benthamiana*. Values were mean ± SD (n = 3). One-way ANOVA followed by post-hoc Tukey’s HSD was applied to evaluate the significance among different heat treatment durations (*p* < 0.05). Samples sharing the same letters showed no significant difference.

To further reveal the regulatory networks of *RtHSFA2a* and *RtHSFA2b* to their targets, we conducted transient effector and promoter-reporter assays in *N. benthamiana* leaves (Wang et al., 2021). We found that *RtHSFA2a* could strongly transactivate *RtHSFA2b* (Figure 10j and n), reinforcing the notion that *RtHSFA2a* transcriptionally activated *RtHSFA2b*. Interestingly, *RtHSFA2b* had significantly higher transactivation activity to *RtHSP21.8* than *RtHSFA2a* (Figure 10k and o); *RtHSFA2b* strongly activated *RtHSP70*, while *RtHSFA2a* transrepressed *RtHSP70* in *N. benthamiana* (Figure 10l and p). All these results indicated that *RtHSFA2a* might function by transactivating *RtHSFA2b*, in turn, *RtHSFA2b* transactivated their targets in *R. tomentosa*.

## 4. Discussion

HSF family might be of great importance in adapting to the hostile terrestrial environment for higher plants (Godfree et al., 2017; Sankoff and Zheng, 2018). Compared with other eukaryotes, plants possess a much larger HSF family. *Drosophila* and *Caenorhabditis elegans* have only one HSF member; yeast has one HSF plus three HSF-related proteins (Nover et al., 2001, 1996), and the human genome encodes six HSF proteins (Gomez-Pastor et al., 2018). Nevertheless, *Arabidopsis* (Nover et al., 2001), common bean (Zhang et al., 2022b), moso bamboo (Huang et al., 2021), poplar (Zhang et al., 2015), and kiwifruit (Ling et al., 2023) possess 21, 30, 41, 28, and 41 members of HSF family respectively. What should be noticed is that lower plants usually have fewer HSF members. There is only one HSF gene in *Volvox carteri*, 6 HSF genes in *Marchantia polymorpha*, and 7 HSF genes in *Selaginella moellendorffii* (Yu et al., 2022). This striking multiplicity provides more regulation flexibility in response to different stages of development or adverse stress, a crucial ability for sessile organisms (Andrási et al., 2021). Among the 25 *HSF* genes we identified in *R. tomentosa*, the class A *RtHSF* occupies the largest proportion and exhibits the highest divergence in evolution, which is consistent with other plants (Liu et al., 2018; Zhang et al., 2020a). The phylogenetic tree reveals the conservation in evolution in the same group shared by many genes. Noteworthy, *R. tomentosa* possesses 2 and 3 members in subclasses A2 and A9 respectively, whilst *Arabidopsis* and *Populus* contain only one member in these two subclasses (Scharf et al., 2012; Zhang et al., 2015). In *Eucalyptus*, a splendid forest tree of *Myrtaceae*, the HSF family has 36 members, and subclass A9 has been enlarged to 16 members (Yuan et al., 2022). We conjectured that *R. tomentosa* and *E. grandis* share similar mechanisms in adapting to tropical climate in evolution and subclasses A2 and A9 may be specialized for this ability.

Gene duplications derived from mutations have long been recognized as contributors of new genes with new functions (Birchler and Yang, 2022). The expansion and distribution of the *RtHSF* family compared with lower plants indicate that *R. tomentosa* has undergone gene duplications in the evolutionary process. We find nine pairs of genes that are the results of gene duplication events, covering classes A, B, and C. Our previous findings reveal that *R. tomentosa* has undergone whole-genome duplication events (Yang et al., 2024). These WGD events, followed by fractionation and neofunctionalization, result in the HSF gene family expansion in *R. tomentosa* (Sankoff and Zheng, 2018). For example, *RtHSFA9a* and *RtHSFA9c* (but not *RtHSFA9b*) are the paralogous genes of *RtHSFA2a* and *RtHSFA2b*. However, *HSFA9* is absent in many species such as kiwifruit (Ling et al., 2023), sesame (Dossa et al., 2016), and *Brassica napus* (Zhu et al., 2017). We speculate that *RtHSFA9a* and *RtHSFA9c* derive from *RtHSFA2* and neofunctionalize thereafter duplication and form a new subclass. The multiplied HSFs and complex modulation of their activities by hetero-oligomerization render complex functions to HSFs in plants (Jacob et al., 2017). HSFB2 and HSFC1 have also undergone expansion and evolution in *R. tomentosa.* As class HSFB and HSFC function as co-regulators of class HSFA (Andrási et al., 2021; Rao et al., 2022; Wu et al., 2024), the enlarged HSFB and HSFC subfamily contribute more flexibility to functional modulation when employing HSFs.

Expression regulation is vital to plants in growth and stress reactions. We identify 25 important cis-acting elements in the promoter regions of *RtHSF* genes according to the results of Plantcare database analysis, and they show large divergences among the *RtHSF* genes. These elements are mainly associated with phytohormone and stress response. For example, ABRE (abscisic acid responsiveness) and DRE (dehydration responsive element) which are related to drought and salt stress response in plants (Narusaka et al., 2003), are presented in the promoters of most HSF genes, while the total number of these two elements is various in each gene, indicating that *RtHSFs* might contribute unequally to drought stress acclimatization. Same with ABRE and DRE, *RtHSF* genes have a high affinity to TGACG-motif (MeJA responsive element) (Zhao et al., 2023), STRE (stress responsive element), and as-1 (activation sequence-1, response to oxidative stress and salicylic acid) (Garretón et al., 2002). These results uphold the performance of HSF genes in stress response including anoxia (Banti et al., 2010), salt (Lü et al., 2022; Zang et al., 2019), drought (Wang et al., 2020; Zhang et al., 2020b), oxidative stress (Satoh et al., 2014), as well as disease (Anckar and Sistonen, 2011; Kumar et al., 2009). Due to the composition of the cis-acting element, the expression profiles of certain *RtHSF* genes exhibit tissue-specific straits. *HSFA9* is expressed specifically in seeds, involved in desiccation tolerance (Prieto-Dapena et al., 2008), seed embryogenesis and early photomorphogenesis (Kotak et al., 2007; Prieto-Dapena et al., 2017), seed longevity (Tejedor-Cano et al., 2010), and UV-B light response (Carranco et al., 2022). The expression level of *RtHSFA9a*, *RtHSFA9b*, and *RtHSFA9c* showed an accumulation in fruit in *R. tomentosa*, consistent with their functions revealed in other plants. *RtHSFB1*, *RtHSFB3*, *RtHSFB5* are highly accumulated in root. Previous research denotes that HSFB1 and HSFB2b are involved in pathogen resistance by regulating Pdf1.2 (Conrath et al., 2015; Kumar et al., 2009), *RtHSFB1* could be presumed that it might be accumulated by the pathogen-rich soil conditions.

We further quantify the mRNA abundance of 9 genes in 4 subclasses under heat stress conditions, and they show significant variance even in the same subclass. Overtly, subclass A2 promptly responds to heat stress in *R. tomentosa*, consistent with previous studies (Gu et al., 2019; Ling et al., 2023; Liu et al., 2023; Wang et al., 2017a); however, we find that *HSFA2b* is induced by heat stress at an exaggerative extent compared with *RtHSFA2a*. We speculate that *RtHSFA2b* functions predominantly in heat stress response. As the *cis-acting* elements vary in the promoter regions of *RtHSFA2a* and *RtHSFA2b*, we wondered whether they had similar differentiation in response to other abiotic stresses. This is further confirmed by the expression profile that *RtHSFA2a* is more sensitive to drought, dehydration, cold, and salt stress, eliciting a notion that *RtHSFA2a* and *RtHSFA2b* have undergone subfunctionalization after gene duplication (Birchler and Yang, 2022).

Considering the expression pattern of *RtHSFA2a* and *RtHSFA2b* under heat stress, we surmise that *RtHSFA2b* is a principal element for *R. tomentosa* thriving in hot and humid tropical climates. Performances of transgenic plants with overexpressed *RtHSFA2a* or *RtHSFA2b* reveal that *RtHSFA2b* plays a pivotal role in handling sudden lethal heat stress rather than *RtHFSA2a*. We then examine the regulatory network of *RtHSFA2a* and *RtHSFA2b* via transiently overexpressing *RtHSFA2a* and *RtHSFA2b* in *R. tomentosa*. According to the results, we find that *RtHSFA2a* plays a role in transcriptionally regulating *RtHSFA2b* (Figure 10 b, j, and n). We further quantify the mRNA abundance of *RtHSP21.3*, *RtHSP21.8*, and *RtHSP26.5*, the homologous genes of which have been testified as downstream genes of HSFA2 in *Populus tomentosa* (Li et al., 2022), as well as *RtHSP17.4* and *RtHSP70*. They are all transcriptionally activated by over-accumulated *RtHSFA2a* and *RtHSFA2b*, reinforcing the conclusions proposed by previous studies (Wang et al., 2017a). What is worth careful deliberation is that, even though most of the quantified HSP genes have higher transcription abundance in OxHSFA2a lines, it could not be concluded that *RtHSFA2a* and *RtHSFA2b* have similar transcription activity. We speculate that *RtHSFA2a* can highly activate *RtHSFA2b* in OxHSFA2a, which in turn activates its target genes. Therefore, the elevated mRNA abundance of HSR genes in the OxHSFA2a line should be regarded as a combination of overaccumulation of *RtHSFA2a* and *RtHSFA2b*, hence *RtHSFA2b* might have higher transcription activity as the variations of HSFA2 in grape (Liu et al., 2023). We verified our conjecture by comparing the transactivation activity of *RtHSFA2a* and *RtHSFA2b* to their targets, and the results prove that *RtHSFA2b* has much higher transcriptional activity to certain targets like *RtHSP21.8* and *RtHSP70* compared with *RtHSFA2a* in *N. benthamiana* (Figure 10). Based on these cognitions, we conclude that *RtHSFA2a* and *RtHSFA2b* all take roles in the thermal adaption mechanisms of *R. tomentosa*, *viz.*, *RtHSFA2a* takes part in the transcriptional activation of *RtHSFA2b*, and *RtHSFA2b*, in turn, dramatically activates its downstream genes to cope with advertise stresses, especially heat stress. However, the regulatory networks of *RtHSFA2a* and *RtSHFA2b* in *R. tomentosa* remain largely unknown, which indubitably requires more elaborate experiments.

Above 3000 species in the *Myrtaceae* family are distributed predominantly in tropical and subtropical regions of the world (Christenhusz and Byng, 2016), indicating that *Myrtaceae* plants possess many advantages a priori in adaption to high temperature and humid conditions. *R.tomentoa*, a representative plant of the *Myrtaceae* family, has captured more and more attention due to its special values. We summarize here that the *R. tomentosa* genome encodes 25 HSF family genes, and classes A, B, and C contain 15, 8, and 2 members respectively. This family has undergone several gene duplications, followed by subfunctionalization or neofunctionalization. Among these 25 HSF genes, *RtHSFA2a* and *RtHSFA2b* show different expression patterns to several abiotic stresses. *RtHSFA2b* is specialized for heat stress for it responds to high temperature more acutely and has higher transcriptional activity to certain HSR genes compared with *RtHSFA2a*, while *RtHSFA2a* functions by transactivating *RtHSFA2b.* This study poses a view of the HSF family at a genome-wide level and reveals a basic thermal adaption mechanism of *R. tomentosa*, a species, we think, can serve as an eminent model plant for adversity.

## Acknowledgments

This research was supported by a grant from the Funding by Science and Technology Projects in Guangzhou (E33309).

## Author contributions

S.D. conceived and supervised the project. H.-G.L. conceived part of the project. H.-G.L. carried out most of the experiments. L.Y. conducted the RNA-Seq of tissues and helped carry out the luciferase assay. L.Y., Y.F., and G.W. helped to prepare plant materials, carry out the stress treatments, and acquire data. S.L. gave invaluable input to the project. S.D. and S.L. acquired funding. H.-G.L. and S.D. analyzed and interpreted the data. H.-G.L. drafted the original manuscript. S.D. reviewed the manuscript. All authors have read and agreed to the published version of the manuscript.

## References

Anckar, J., Sistonen, L., 2011. Regulation of HSF1 function in the heat stress response: implications in aging and disease. Annu. Rev. Biochem. 80, 1089–1115.

Andr ási, N., Pettkó-Szandtner, A., Szabados, L., 2021. Diversity of plant heat shock factors: regulation, interactions, and functions. J. Exp. Bot. 72, 1558–1575.

Baniwal, S.K., Chan, K.Y., Scharf, K.D., Nover, L., 2007. Role of heat stress transcription factor HsfA5 as specific repressor of HsfA4. J. Biol. Chem. 282, 3605–3613.

Banti, V., Mafessoni, F., Loreti, E., Alpi, A., Perata, P., 2010. The heat-inducible transcription factor *HsfA2* enhances anoxia tolerance in *Arabidopsis*. Plant Physiol. 152, 1471–1483.

Bhat, J.A., Kundu, M.C., Hazra, G.C., Santra, G.H., Mandal, B., 2010. Rehabilitating acid soils for increasing crop productivity through low-cost liming material. Sci. Total Environ. 408, 4346–4353.

Biggin, A.J., Piispa, E.J., Pesonen, L.J., Holme, R., Paterson, G.A., Veikkolainen, T., Tauxe, L., 2015. Palaeomagnetic field intensity variations suggest Mesoproterozoic inner-core nucleation. Nature 526, 245– 248.

Birchler, J.A., Yang, H., 2022. The multiple fates of gene duplications: Deletion, hypofunctionalization, subfunctionalization, neofunctionalization, dosage balance constraints, and neutral variation. Plant Cell 34, 2466–2474.

Carranco, R., Prieto-Dapena, P., Almoguera, C., Jordano, J., 2022. A seed-specific transcription factor, HSFA9, anticipates UV-B light responses by mimicking the activation of the UV-B receptor in tobacco. Plant J. 111, 1439–1452.

Chen, C., Wu, Y., Li, J., Wang, X., Zeng, Z., Xu, J., Liu, Y., Feng, J., Chen, H., He, Y., Xia, R., 2023a. TBtools-II: A “one for all, all for one” bioinformatics platform for biological big-data mining. Mol. Plant 16, 1733–1742.

Chen, H., Liu, X., Li, S., Yuan, L., Mu, H., Wang, Y., Li, Y., Duan, W., Fan, P., Liang, Z., Wang, L., 2023b. The class B heat shock factor HSFB1 regulates heat tolerance in grapevine. Hortic. Res. 10, uhad001.

Christenhusz, M.J.M., Byng, J.W., 2016. The number of known plants species in the world and its annual increase. Phytotaxa 261, 201–217.

Clough, S.J., Bent, A.F., 1998. Floral dip: a simplified method for *Agrobacterium*-mediated transformation of *Arabidopsis thaliana*. Plant J. 16, 735–743.

Conrath, U., Beckers, G.J.M., Langenbach, C.J.G., Jaskiewicz, M.R., 2015. Priming for enhanced defense. Annu. Rev. Phytopathol. 53, 97–119.

Doglioni, C., Pignatti, J., Coleman, M., 2016. Why did life develop on the surface of the Earth in the Cambrian? Geosci. Front. 7, 865–873.

Dossa, K., Diouf, D., Cisse, N., 2016. Genome-wide investigation of Hsf genes in sesame reveals their segmental duplication expansion and their active role in drought stress response. Front. Plant Sci. 7, 1522.

Flores, B.M., Montoya, E., Sakschewski, B., Nascimento, N., Staal, A., Betts, R.A., Levis, C., Lapola, D.M., Esquí vel-Muelbert, A., Jakovac, C., Nobre, C.A., Oliveira, R.S., Borma, L.S., Nian, D., Boers, N., Hecht, S.B., Ter Steege, H., Arieira, J., Lucas, I.L., Berenguer, E., Marengo, J.A., Gatti, L.V., Mattos, C.R.C., Hirota, M., 2024. Critical transitions in the Amazon forest system. Nature 626, 555–564.

Friedrich, T., Oberkofler, V., Trindade, I., Altmann, S., Brzezinka, K., Lämke, J., Gorka, M., Kappel, C., Sokolowska, E., Skirycz, A., Graf, A., Bäurle, I., 2021. Heteromeric HSFA2/HSFA3 complexes drive transcriptional memory after heat stress in *Arabidopsis*. Nat. Commun. 12, 3426.

Garretón, V., Carpinelli, J., Jordana, X., Holuigue, L., 2002. The *as-1* promoter element is an oxidative stress-responsive element and salicylic acid activates it via oxidative species. Plant Physiol. 130, 1516– 1526.

Godfree, R.C., Marshall, D.J., Young, A.G., Miller, C.H., Mathews, S., 2017. Empirical evidence of fixed and homeostatic patterns of polyploid advantage in a keystone grass exposed to drought and heat stress. R. Soc. Open Sci. 4, 170934.

Gomez-Pastor, R., Burchfiel, E.T., Thiele, D.J., 2018. Regulation of heat shock transcription factors and their roles in physiology and disease. Nat. Rev., Mol. Cell Biol. 19, 4–19.

Goodstein, D.M., Shu, S., Howson, R., Neupane, R., Hayes, R.D., Fazo, J., Mitros, T., Dirks, W., Hellsten, U., Putnam, N., Rokhsar, D.S., 2012. Phytozome: a comparative platform for green plant genomics. Nucleic Acids Res. 40, D1178–D1186.

Gu, L., Jiang, T., Zhang, C., Li, X., Wang, C., Zhang, Y., Li, T., Dirk, L.M.A., Downie, A.B., Zhao, T., 2019. Maize HSFA2 and HSBP2 antagonistically modulate raffinose biosynthesis and heat tolerance in *Arabidopsis*. Plant J. 100, 128–142.

Guo, M., Liu, J.H., Ma, X., Luo, D.X., Gong, Z.H., Lu, M.H., 2016. The plant heat stress transcription factors (HSFs): Structure, regulation, and function in response to abiotic stresses. Front. Plant Sci. 7, 114.

Harrison, C.J., Bohm, A.A., Nelson, H.C., 1994. Crystal structure of the DNA binding domain of the heat shock transcription factor. Science 263, 224–227.

He, S.-M., Wang, X., Yang, S.-C., Dong, Y., Zhao, Q.-M., Yang, J.-L., Cong, K., Zhang, J.-J., Zhang, G.-H., Wang, Y., Fan, W., 2018. *De novo* transcriptome characterization of Rhodomyrtus tomentosa leaves and identification of genes involved in α/β-pinene and β-caryophyllene biosynthesis. Front. Plant Sci. 9, 1231.

Huang, B., Huang, Z., Ma, R., Chen, J., Zhang, Z., Yrjälä, K., 2021. Genome-wide identification and analysis of the heat shock transcription factor family in moso bamboo (*Phyllostachys edulis*). Sci. Rep. 11, 16492.

Ikeda, M., Mitsuda, N., Ohme-Takagi, M., 2011. *Arabidopsis* HsfB1 and HsfB2b act as repressors of the expression of heat-inducible *Hsfs* but positively regulate the acquired thermotolerance. Plant Physiol. 157, 1243–1254.

Jacob, P., Hirt, H., Bendahmane, A., 2017. The heat-shock protein/chaperone network and multiple stress resistance. Plant Biotechnol. J. 15, 405–414.

Jiao, Y., Wickett, N.J., Ayyampalayam, S., Chanderbali, A.S., Landherr, L., Ralph, P.E., Tomsho, L.P., Hu, Y., Liang, H., Soltis, P.S., Soltis, D.E., Clifton, S.W., Schlarbaum, S.E., Schuster, S.C., Ma, H., Leebens-Mack, J., dePamphilis, C.W., 2011. Ancestral polyploidy in seed plants and angiosperms. Nature 473, 97–100.

Kan, Y., Mu, X.-R., Gao, J., Lin, H.-X., Lin, Y., 2023. The molecular basis of heat stress responses in plants. Mol. Plant 16, 1612–1634.

Kan, Y., Mu, X.-R., Zhang, H., Gao, J., Shan, J.-X., Ye, W.-W., Lin, H.-X., 2021. *TT2* controls rice thermotolerance through SCT1-dependent alteration of wax biosynthesis. Nat. Plants 8, 53–67.

Kanwar, M., Chaudhary, C., Anand, K.A., Singh, S., Garg, M., Mishra, S.K., Sirohi, P., Chauhan, H., 2023. An insight into *Pisum sativum* HSF gene family-Genome-wide identification, phylogenetic, expression, and analysis of transactivation potential of pea heat shock transcription factor. Plant Physiol. Biochem. 202, 107971.

Kenrick, P., Crane, P.R., 1997. The origin and early evolution of plants on land. Nature 389, 33–39.

Kochian, L.V., Piñeros, M.A., Liu, J., Magalhaes, J.V., 2015. Plant adaptation to acid soils: The molecular basis for crop aluminum resistance. Annu. Rev. Plant Biol. 66, 571–598.

Kotak, S., Port, M., Ganguli, A., Bicker, F., von Koskull-Doring, P., 2004. Characterization of C-terminal domains of *Arabidopsis* heat stress transcription factors (Hsfs) and identification of a new signature combination of plant class A Hsfs with AHA and NES motifs essential for activator function and intracellular localization. Plant J. 39, 98–112.

Kotak, S., Vierling, E., Bäumlein, H., von Koskull-Döring, P., 2007. A novel transcriptional cascade regulating expression of heat stress proteins during seed development of *Arabidopsis*. Plant Cell 19, 182–195.

Kozlov, A.M., Darriba, D., Flouri, T., Morel, B., Stamatakis, A., 2019. RAxML-NG: a fast, scalable and user-friendly tool for maximum likelihood phylogenetic inference. Bioinformatics 35, 4453–4455.

Kumar, M., Busch, W., Birke, H., Kemmerling, B., Nürnberger, T., Schöffl, F., 2009. Heat shock Factors HsfB1 and HsfB2b are involved in the regulation of Pdf1.2 expression and pathogen resistance in *Arabidopsis*. Mol. Plant 2, 152–165.

Lai, T.N.H., André, C., Rogez, H., Mignolet, E., Nguyen, T.B.T., Larondelle, Y., 2015. Nutritional composition and antioxidant properties of the sim fruit (*Rhodomyrtus tomentosa*). Food Chem. 168, 410–416.

Lang, S., Liu, X., Xue, H., Li, X., Wang, X., 2017. Functional characterization of BnHSFA4a as a heat shock transcription factor in controlling the re-establishment of desiccation tolerance in seeds. J. Exp. Bot. 68, 2361–2375.

Li, H.-G., Yang, Y., Liu, M., Zhu, Y., Wang, H.-L., Feng, C.-H., Niu, M.-X., Liu, C., Yin, W., Xia, X., 2022. The in vivo performance of a heat shock transcription factor from *Populus euphratica, PeHSFA2*, promises a prospective strategy to alleviate heat stress damage in poplar. Environ. Exp. Bot. 201, 104940.

Li, X.-L., Meng, D., Li, M.-J., Zhou, J., Yang, Y.-Z., Zhou, B.-B., Wei, Q.-P., Zhang, J.-K., 2023. Transcription factors MhDREB2A/MhZAT10 play a role in drought and cold stress response crosstalk in apple. Plant Physiol. 192, 2203–2220.

Li, Y., Chen, T., Wang, W., Liu, H., Yan, X., Wu-Zhang, K., Qin, W., Xie, L., Zhang, Y., Peng, B., Yao, X., Wang, C., Kayani, S.-I., Fu, X., Li, L., Tang, K., 2021. A high-efficiency *Agrobacterium*-mediated transient expression system in the leaves of *Artemisia annua* L. Plant Methods 17, 106.

Li, Z., Li, Z., Ji, Y., Wang, C., Wang, S., Shi, Y., Le, J., Zhang, M., 2024. The Heat shock factor 20-HSF4-Cellulose synthase A2 module regulates heat stress tolerance in maize. Plant Cell koae106.

Ling, C., Liu, Y., Yang, Z., Xu, J., Ouyang, Z., Yang, J., Wang, S., 2023. Genome-wide identification of HSF gene family in kiwifruit and the function of *AeHSFA2b* in salt tolerance. Int. J. Mol. Sci. 24, 15638.

Liu, G., Chai, F., Wang, Y., Jiang, J., Duan, W., Wang, Y., Wang, F., Li, S., Wang, L., 2018. Genome-wide identification and classification of HSF family in grape, and their transcriptional analysis under heat acclimation and heat stress. Hortic. Plant J. 4, 133–143.

Liu, H.-C., Liao, H.-T., Charng, Y.-Y., 2011. The role of class A1 heat shock factors (HSFA1s) in response to heat and other stresses in *Arabidopsis*. Plant Cell Environ. 34, 738–751.

Liu, X., Chen, H., Li, S., Lecourieux, D., Duan, W., Fan, P., Liang, Z., Wang, L., 2023. Natural variations of HSFA2 enhance thermotolerance in grapevine. Hortic. Res. 10, uhac250.

Livak, K.J., Schmittgen, T.D., 2001. Analysis of relative gene expression data using real-time quantitative PCR and the 2^−ΔΔCT^ method. Methods 25, 402–408.

Lobell, D.B., Schlenker, W., Costa-Roberts, J., 2011. Climate trends and global crop production since 1980. Science 333, 616–620.

Lü, X.-P., Shao, K.-Z., Xu, J.-Y., Li, J.-L., Ren, W., Chen, J., Zhao, L.-Y., Zhao, Q., Zhang, J.-L., 2022. A heat shock transcription factor gene (*HaHSFA1*) from a desert shrub, *Haloxylon ammodendron*, elevates salt tolerance in *Arabidopsis thaliana*. Environ. Exp. Bot. 201, 104954.

Mittal, D., Chakrabarti, S., Sarkar, A., Singh, A., Grover, A., 2009. Heat shock factor gene family in rice: Genomic organization and transcript expression profiling in response to high temperature, low temperature and oxidative stresses. Plant Physiol. Biochem. 47, 785–795.

Mo, L., Zohner, C.M., Reich, P.B., Liang, J., de Miguel, S., Nabuurs, G.-J., Renner, S.S., van den Hoogen, J., Araza, A., Herold, M., Mirzagholi, L., Ma, H., Averill, C., Phillips, O.L., Gamarra, J.G.P., Hordijk, I., Routh, D., Abegg, M., Adou Yao, Y.C., Alberti, G., Almeyda Zambrano, A.M., Alvarado, B.V., Alvarez-Dávila, E., Alvarez-Loayza, P., Alves, L.F., Amaral, I., Ammer, C., Antón-Fernández, C., Araujo-Murakami, A., Arroyo, L., Avitabile, V., Aymard, G.A., Baker, T.R., Bałazy, R., Banki, O., Barroso, J.G., Bastian, M.L., Bastin, J.-F., Birigazzi, L., Birnbaum, P., Bitariho, R., Boeckx, P., Bongers, F., Bouriaud, O., Brancalion, P.H.S., Brandl, S., Brearley, F.Q., Brienen, R., Broadbent, E.N., Bruelheide, H., Bussotti, F., Cazzolla Gatti, R., César, R.G., Cesljar, G., Chazdon, R.L., Chen, H.Y.H., Chisholm, C., Cho, H., Cienciala, E., Clark, C., Clark, D., Colletta, G.D., Coomes, D.A., Cornejo Valverde, F., Corral-Rivas, J.J., Crim, P.M., Cumming, J.R., Dayanandan, S., de Gasper, A.L., Decuyper, M., Derroire, G., DeVries, B., Djordjevic, I., Dolezal, J., Dourdain, A., Engone Obiang, N.L., Enquist, B.J., Eyre, T.J., Fandohan, A.B., Fayle, T.M., Feldpausch, T.R., Ferreira, L.V., Finér, L., Fischer, M., Fletcher, C., Frizzera, L., Gianelle, D., Glick, H.B., Harris, D.J., Hector, A., Hemp, A., Hengeveld, G., Hérault, B., Herbohn, J.L., Hillers, A., Honorio Coronado, E.N., Hui, C., Ibanez, T., Imai, N., Jagodziński, A.M., Jaroszewicz, B., Johannsen, V.K., Joly, C.A., Jucker, T., Jung, I., Karminov, V., Kartawinata, K., Kearsley, E., Kenfack, D., Kennard, D.K., Kepfer-Rojas, S., Keppel, G., Khan, M.L., Killeen, T.J., Kim, H.S., Kitayama, K., Köhl, M., Korjus, H., Kraxner, F., Kucher, D., Laarmann, D., Lang, M., Lu, H., Lukina, N.V., Maitner, B.S., Malhi, Y., Marcon, E., Marimon, B.S., Marimon-Junior, B.H., Marshall, A.R., Martin, E.H., Meave, J.A., Melo-Cruz, O., Mendoza, C., Mendoza-Polo, I., Miscicki, S., Merow, C., Monteagudo Mendoza, A., Moreno, V.S., Mukul, S.A., Mundhenk, P., Nava-Miranda, M.G., Neill, D., Neldner, V.J., Nevenic, R.V., Ngugi, M.R., Niklaus, P.A., Oleksyn, J., Ontikov, P., Ortiz-Malavasi, E., Pan, Y., Paquette, A., Parada-Gutierrez, A., Parfenova, E.I., Park, M., Parren, M., Parthasarathy, N., Peri, P.L., Pfautsch, S., Picard, N., Piedade, M.T.F., Piotto, D., Pitman, N.C.A., Poulsen, A.D., Poulsen, J.R., Pretzsch, H., Ramirez Arevalo, F., Restrepo-Correa, Z., Rodeghiero, M., Rolim, S.G., Roopsind, A., Rovero, F., Rutishauser, E., Saikia, P., Salas-Eljatib, C., Saner, P., Schall, P., Schelhaas, M.-J., Schepaschenko, D., Scherer-Lorenzen, M., Schmid, B., Schöngart, J., Searle, E.B., Seben, V., Serra-Diaz, J.M., Sheil, D., Shvidenko, A.Z., Silva-Espejo, J.E., Silveira, M., Singh, J., Sist, P., Slik, F., Sonké, B., Souza, A.F., Stereńczak, K.J., Svenning, J.-C., Svoboda, M., Swanepoel, B., Targhetta, N., Tchebakova, N., Ter Steege, H., Thomas, R., Tikhonova, E., Umunay, P.M., Usoltsev, V.A., Valencia, R., Valladares, F., van der Plas, F., Van Do, T., van Nuland, M.E., Vasquez, R.M., Verbeeck, H., Viana, H., Vibrans, A.C., Vieira, S., von Gadow, K., Wang, H.-F., Watson, J.V., Werner, G.D.A., Wiser, S.K., Wittmann, F., Woell, H., Wortel, V., Zagt, R., Zawiła-Niedźwiecki, T., Zhang, C., Zhao, X., Zhou, M., Zhu, Z.-X., Zo-Bi, I.C., Gann, G.D., Crowther, T.W., 2023. Integrated global assessment of the natural forest carbon potential. Nature 624, 92–101.

Narusaka, Y., Nakashima, K., Shinwari, Z.K., Sakuma, Y., Furihata, T., Abe, H., Narusaka, M., Shinozaki, K., Yamaguchi-Shinozaki, K., 2003. Interaction between two cis-acting elements, ABRE and DRE, in ABA-dependent expression of *Arabidopsis r*d29A gene in response to dehydration and high-salinity stresses. Plant J. 34, 137–148.

Nover, L., Bharti, K., Döring, P., Mishra, S.K., Ganguli, A., Scharf, K.D., 2001. *Arabidopsis* and the heat stress transcription factor world: how many heat stress transcription factors do we need? Cell Stress Chaperon. 6, 177–189.

Nover, L., Scharf, K.-D., Gagliardi, D., Vergne, P., Czarnecka-Verner, E., Gurley, W.B., 1996. The Hsf world: classification and properties of plant heat stress transcription factors. Cell Stress Chaperon. 1, 215–223.

Ohama, N., Sato, H., Shinozaki, K., Yamaguchi-Shinozaki, K., 2017. Transcriptional regulatory network of plant heat stress response. Trends Plant Sci. 22, 53–65.

Pecl, G.T., Araújo, M.B., Bell, J.D., Blanchard, J., Bonebrake, T.C., Chen, I.-C., Clark, T.D., Colwell, R.K., Danielsen, F., Evengård, B., Falconi, L., Ferrier, S., Frusher, S., Garcia, R.A., Griffis, R.B., Hobday, A.J., Janion-Scheepers, C., Jarzyna, M.A., Jennings, S., Lenoir, J., Linnetved, H.I., Martin, V.Y., McCormack, P.C., McDonald, J., Mitchell, N.J., Mustonen, T., Pandolfi, J.M., Pettorelli, N., Popova, E., Robinson, S.A., Scheffers, B.R., Shaw, J.D., Sorte, C.J.B., Strugnell, J.M., Sunday, J.M., Tuanmu, M.-N., Vergés, A., Villanueva, C., Wernberg, T., Wapstra, E., Williams, S.E., 2017. Biodiversity redistribution under climate change: Impacts on ecosystems and human well-being. Science 355, eaai9214.

Prieto-Dapena, P., Almoguera, C., Personat, J.-M., Merchan, F., Jordano, J., 2017. Seed-specific transcription factor HSFA9 links late embryogenesis and early photomorphogenesis. J. Exp. Bot. 68, 1097–1108.

Prieto-Dapena, P., Castano, R., Almoguera, C., Jordano, J., 2008. The ectopic overexpression of a seed-specific transcription factor, HaHSFA9, confers tolerance to severe dehydration in vegetative organs. Plant J. 54, 1004–1014.

Qiao, X., Li, Q., Yin, H., Qi, K., Li, L., Wang, R., Zhang, S., Paterson, A.H., 2019. Gene duplication and evolution in recurring polyploidization-diploidization cycles in plants. Genome Biol 20, 38.

Qin, W., Wang, N., Yin, Q., Li, H., Wu, A.-M., Qin, G., 2022. Activation tagging identifies WRKY14 as a repressor of plant thermomorphogenesis in *Arabidopsis*. Mol. Plant 15, 1725–1743.

Rao, S., Das, J.R., Balyan, S., Verma, R., Mathur, S., 2022. Cultivar-biased regulation of *HSFA7* and *HSFB4a* govern high-temperature tolerance in tomato. Planta 255, 31.

Sankoff, D., Zheng, C., 2018. Whole genome duplication in plants: Implications for evolutionary analysis. Methods Mol. Biol. 1704, 291–315.

Satoh, M., Tokaji, Y., Nagano, A.J., Hara-Nishimura, I., Hayashi, M., Nishimura, M., Ohta, H., Masuda, S., 2014. *Arabidopsis* mutants affecting oxylipin signaling in photo-oxidative stress responses. Plant Physiol. Biochem. 81, 90–95.

Scharf, K.-D., Berberich, T., Ebersberger, I., Nover, L., 2012. The plant heat stress transcription factor (Hsf) family: structure, function and evolution. Biochim. Biophys. Acta 1819, 104–119.

Schreiber, M., Rensing, S.A., Gould, S.B., 2022. The greening ashore. Trends Plant Sci. 27, 847–857.

Sinaga, E., Suprihatin, Yenisbar, Iswahyudi, M., Setyowati, S., Prasasty, V.D., 2021. Effect of supplementation of *Rhodomyrtus tomentosa* fruit juice in preventing hypercholesterolemia and atherosclerosis development in rats fed with high fat high cholesterol diet. Biomed. Pharmacother. 142, 111996.

Srisuwan, S., Mackin, K.E., Hocking, D., Lyras, D., Bennett-Wood, V., Voravuthikunchai, S.P., Robins-Browne, R.M., 2018. Antibacterial activity of rhodomyrtone on *Clostridium difficile* vegetative cells and spores in vitro. Int. J. Antimicrob. Agents 52, 724–729.

Tejedor-Cano, J., Prieto-Dapena, P., Almoguera, C., Carranco, R., Hiratsu, K., Ohme-Takagi, M., Jordano, J., 2010. Loss of function of the HSFA9 seed longevity program. Plant Cell Environ. 33, 1408–1417.

Wang, H., Feng, M., Jiang, Y., Du, D., Dong, C., Zhang, Z., Wang, W., Liu, J., Liu, X., Li, S., Chen, Y., Guo, W., Xin, M., Yao, Y., Ni, Z., Sun, Q., Peng, H., Liu, J., 2023. Thermosensitive SUMOylation of TaHsfA1 defines a dynamic ON/OFF molecular switch for the heat stress response in wheat. Plant Cell 35, 3889–3910.

Wang, H.-L., Zhang, Y., Wang, T., Yang, Q., Yang, Y., Li, Z., Li, B., Wen, X., Li, W., Yin, W., Xia, X., Guo, H., Li, Z., 2021. An alternative splicing variant of PtRD26 delays leaf senescence by regulating multiple NAC transcription factors in *Populus*. Plant Cell 33, 1594–1614.

Wang, N., Liu, W., Yu, L., Guo, Z., Chen, Z., Jiang, S., Xu, H., Fang, H., Wang, Y., Zhang, Z., Chen, X., 2020. HEAT SHOCK FACTOR A8a modulates flavonoid synthesis and drought tolerance. Plant Physiol. 184, 1273–1290.

Wang, R., Yao, L., Lin, X., Hu, X., Wang, L., 2022a. Exploring the potential mechanism of *Rhodomyrtus tomentosa* (Ait.) Hassk fruit phenolic rich extract on ameliorating nonalcoholic fatty liver disease by integration of transcriptomics and metabolomics profiling. Food Res. Int. 151, 110824.

Wang, R., Yao, L., Meng, T., Li, C., Wang, L., 2022b. *Rhodomyrtus tomentosa* (Ait.) Hassk fruit phenolic-rich extract mitigates intestinal barrier dysfunction and inflammation in mice. Food Chem. 393, 133438.

Wang, X., Huang, W., Liu, J., Yang, Z., Huang, B., 2017a. Molecular regulation and physiological functions of a novel *FaHsfA2c* cloned from tall fescue conferring plant tolerance to heat stress. Plant Biotechnol. J. 15, 237–248.

Wang, Y., Pan, F., Chen, D., Chu, W., Liu, H., Xiang, Y., 2017b. Genome-wide identification and analysis of the *Populus trichocarpa* TIFY gene family. Plant Physiol. Biochem. 115, 360–371.

Wang, Y., Tang, H., Debarry, J.D., Tan, X., Li, J., Wang, X., Lee, T., Jin, H., Marler, B., Guo, H., Kissinger, J.C., Paterson, A.H., 2012. *MCScanX*: a toolkit for detection and evolutionary analysis of gene synteny and collinearity. Nucleic Acids Res. 40, e49.

White, E.J., Venter, M., Hiten, N.F., Burger, J.T., 2008. Modified Cetyltrimethylammonium bromide method improves robustness and versatility: The benchmark for plant RNA extraction. Biotechnol. J. 3, 1424– 1428.

Wu, Z., Li, T., Ding, L., Wang, C., Teng, R., Xu, S., Cao, X., Teng, N., 2024. Lily LlHSFC2 coordinates with HSFAs to balance heat stress response and improve thermotolerance. New Phytol. 241, 2124–2142.

Xie, K., Guo, J., Wang, S., Ye, W., Sun, F., Zhang, C., Xi, Y., 2023. Genome-wide identification, classification, and expression analysis of heat shock transcription factor family in switchgrass (*Panicum virgatum* L.). Plant Physiol. Biochem. 201, 107848.

Yang, F.-S., Nie, S., Liu, H., Shi, T.-L., Tian, X.-C., Zhou, S.-S., Bao, Y.-T., Jia, K.-H., Guo, J.-F., Zhao, W., An, N., Zhang, R.-G., Yun, Q.-Z., Wang, X.-Z., Mannapperuma, C., Porth, I., El-Kassaby, Y.A., Street, N.R., Wang, X.-R., Van de Peer, Y., Mao, J.-F., 2020. Chromosome-level genome assembly of a parent species of widely cultivated azaleas. Nat. Commun. 11, 5269.

Yang, L., Jin, J., Lyu, S., Zhang, F., Cao, P., Qin, Q., Zhang, G., Feng, C., Lu, P., Li, H., Deng, S., 2024. Genomic analysis based on chromosome-level genome assembly reveals Myrtaceae evolution and terpene biosynthesis of rose myrtle. BMC genom. 25, 578.

Yang, L., Wu, K., Qiu, S., Cao, H., Deng, S., 2023. A plant regeneration protocol from callus cultures of medicinal plant *Rhodomyrtus tomentosa*. Plant Cell Tiss. Org. 153, 307–317.

Yu, T., Bai, Y., Liu, Z., Wang, Z., Yang, Q., Wu, T., Feng, S., Zhang, Y., Shen, S., Li, Q., Gu, L., Song, X., 2022. Large-scale analyses of heat shock transcription factors and database construction based on whole-genome genes in horticultural and representative plants. Hortic. Res. 9, uhac035.

Yuan, T., Liang, J., Dai, J., Zhou, X.-R., Liao, W., Guo, M., Aslam, M., Li, S., Cao, G., Cao, S., 2022. Genome-wide identification of *Eucalyptus* heat shock transcription factor family and their transcriptional analysis under salt and temperature stresses. Int. J. Mol. Sci. 23, 8044.

Zandalinas, S.I., Fritschi, F.B., Mittler, R., 2021. Global warming, climate change, and environmental pollution: Recipe for a multifactorial stress combination disaster. Trends Plant Sci. 26, 588–599.

Zang, D., Wang, J., Zhang, X., Liu, Z., Wang, Y., 2019. *Arabidopsis* heat shock transcription factor HSFA7b positively mediates salt stress tolerance by binding to an E-box-like motif to regulate gene expression. J. Exp. Bot. 70, 5355–5374.

Zhang, H., Li, G., Fu, C., Duan, S., Hu, D., Guo, X., 2020a. Genome-wide identification, transcriptome analysis and alternative splicing events of Hsf family genes in maize. Sci. Rep. 10, 8073.

Zhang, H., Xiang, Y., He, N., Liu, X., Liu, H., Fang, L., Zhang, F., Sun, X., Zhang, D., Li, X., Terzaghi, W., Yan, J., Dai, M., 2020b. Enhanced vitamin C production mediated by an ABA-induced PTP-like nucleotidase improves plant drought tolerance in *Arabidopsis* and maize. Mol. Plant 13, 760–776.

Zhang, H., Zhou, J.-F., Kan, Y., Shan, J.-X., Ye, W.-W., Dong, N.-Q., Guo, T., Xiang, Y.-H., Yang, Y.-B., Li, Y.-C., Zhao, H.-Y., Yu, H.-X., Lu, Z.-Q., Guo, S.-Q., Lei, J.-J., Liao, B., Mu, X.-R., Cao, Y.-J., Yu, J.-J., Lin, Y., Lin, H.-X., 2022a. A genetic module at one locus in rice protects chloroplasts to enhance thermotolerance. Science 376, 1293–1300.

Zhang, J., Li, X.-M., Lin, H.-X., Chong, K., 2019. Crop improvement through temperature resilience. Annu. Rev. Plant Biol. 70, 753–780.

Zhang, J., Liu, B., Li, J., Zhang, L., Wang, Y., Zheng, H., Lu, M., Chen, J., 2015. *Hsf* and *Hsp* gene families in *Populus*: genome-wide identification, organization and correlated expression during development and in stress responses. BMC genom. 16, 181.

Zhang, Q., Geng, J., Du, Y., Zhao, Q., Zhang, W., Fang, Q., Yin, Z., Li, J., Yuan, X., Fan, Y., Cheng, X., Du, J., 2022b. Heat shock transcription factor (Hsf) gene family in common bean (*Phaseolus vulgaris*): genome-wide identification, phylogeny, evolutionary expansion and expression analyses at the sprout stage under abiotic stress. BMC plant biol. 22, 33.

Zhao, Z., Wu, L., Xie, J., Feng, Y., Tian, J., He, X., Li, B., Wang, L., Wang, X., Zhang, Y., Wu, S., Zheng, X., 2020. *Rhodomyrtus tomentosa* (Aiton.): A review of phytochemistry, pharmacology and industrial applications research progress. Food Chem. 309, 125715.

Zhao, Z., Zhang, R., Wang, D., Zhang, J., Zang, S., Zou, W., Feng, A., You, C., Su, Y., Wu, Q., Que, Y., 2023. Dissecting the features of TGA gene family in *Saccharum* and the functions of *ScTGA1* under biotic stresses. Plant physiol. biochem. 200, 107760.

Zheng, L., Yang, J., Chen, Y., Ding, L., Wei, J., Wang, H., 2021. An improved and efficient method of *Agrobacterium syringe* infiltration for transient transformation and its application in the elucidation of gene function in poplar. BMC plant biol. 21, 54.

Zhu, X., Huang, C., Zhang, L., Liu, H., Yu, J., Hu, Z., Hua, W., 2017. Systematic analysis of Hsf family genes in the *Brassica napus* genome reveals novel responses to heat, drought and high CO_2_ stresses. Front. Plant Sci. 8, 1174.

Zhuang, L., Chen, L.-F., Zhang, Y.-B., Liu, Z., Xiao, X.-H., Tang, W., Wang, G.-C., Song, W.-J., Li, Y.-L., Li, M.-M., 2017. Watsonianone A from *Rhodomyrtus tomentosa* fruit attenuates respiratory-syncytial-virus-induced inflammation in vitro. J. Agric. Food Chem. 65, 3481–3489.

